# Dynamic Metal–Metal Distance Modulation Controls Oxygen Activation in Non-heme Diiron Enzymes

**DOI:** 10.64898/2026.02.02.702239

**Authors:** Simran Nisha, Arunima Choudhury, Subhendu Roy

## Abstract

Multinuclear metal centers catalyze some of the most challenging oxidative transformations in biological and chemical catalysis, yet the molecular principles controlling oxygen activation in non-heme diiron enzymes remain unclear. Structural studies of desaturase-like diiron enzymes frequently reveal elongated Fe–Fe separations lacking canonical bridging motifs, whereas spectroscopic measurements indicate substantial metal–metal coupling during catalysis, creating a longstanding mechanistic paradox between structural and spectroscopic observations. Here we show that oxygen activation can be dynamically regulated by fluctuations in metal–metal distance through extensive multiscale simulations of the membrane-bound fatty-acid decarboxylase UndB. UndB catalyses the conversion of naturally abundant free fatty acids into terminal 1-alkenes–valuable hydrocarbon precursors for sustainable biofuels, and exhibits unusual hydrogen peroxide formation that has led to conflicting mechanistic proposals. Combining quantum mechanics/molecular mechanics (QM/MM) simulations with free energy analysis and quantum chemical calculations, we identify a catalytically competent ensemble in which transient Fe–Fe contraction establishes effective metal–metal coupling and enables formation of a peroxodiiron(III/III) intermediate despite elongated resting-state structures and the absence of canonical bridging motifs. The computed free-energy landscape quantitatively reproduces experimentally inferred catalytic barriers and accounts for the substantial H_2_O_2_ formation observed during catalysis. The reactive state resembles a P-like intermediate rather than the canonical diamond-core Q species commonly invoked for related diiron enzymes. Comparison with magnetic coupling and mechanistic features reported for related non-heme diiron enzymes, including Alkane monooxygenase AlkB, Stearoyl-CoA Desaturase 1 (SCD1) and the soluble decarboxylase UndA, suggests that dynamic metal–metal distance modulation represents a general mechanism governing oxygen activation across diverse diiron enzyme families. These findings establish metal–metal distance modulation as a previously unrecognized control parameter for oxygen activation in diiron and related multinuclear metalloenzymes, reconciling structural and spectroscopic observations and revealing how protein conformational dynamics regulate electronic coupling to control oxidative catalysis, thereby suggesting a general principle for tuning reactivity in multinuclear catalysts.

## Introduction

Oxygen activation by non-heme diiron enzymes (NHFe_2_) remains a long-standing mechanistic paradox^1-3^. Spectroscopic studies consistently indicate substantial metal–metal coupling during catalysis, yet most available structures display elongated Fe–Fe separations that appear incompatible with these observations^4^. This apparent discrepancy between structural and spectroscopic evidence has persisted across desaturase-like, histidine-rich diiron enzyme families, leaving the molecular basis of oxygen activation unresolved. Non-heme diiron centres are widespread in biological redox chemistry and occur in diverse enzymes, including Methane Monooxygenase, Ribonucleotide Reductase and Δ9-Desaturase, where coupling between the two iron centres plays a central role in oxygen activation and substrate oxidation^5^. In contrast, the catalytic mechanisms of recently discovered membrane-bound desaturase-like NHFe_2_ enzymes remain poorly understood, and differing mechanistic models have been proposed to explain their reactivity. Here, we focus on the membrane-bound fatty-acid decarboxylase UndB^3^ which catalyzes oxidative decarboxylation of fatty acids to generate terminal 1-alkenes–valuable hydrocarbon precursors for sustainable biofuels. Unlike typical fatty-acid desaturases, UndB performs oxidative C–C bond cleavage and exhibits high catalytic turnover compared with other fatty acid decarboxylases, including the heme-containing cytochrome P450 enzyme OleT^6^ and the soluble non-heme diiron enzyme UndA^7^. UndB has attracted considerable interest because it catalyzes the selective conversion of diverse fatty acids into terminal alkenes through activation of strong C–H bonds (bond dissociation energy > 95 kcal/mol)^8^, a chemically challenging transformation (Fig. 1a).

**Fig. 1.**
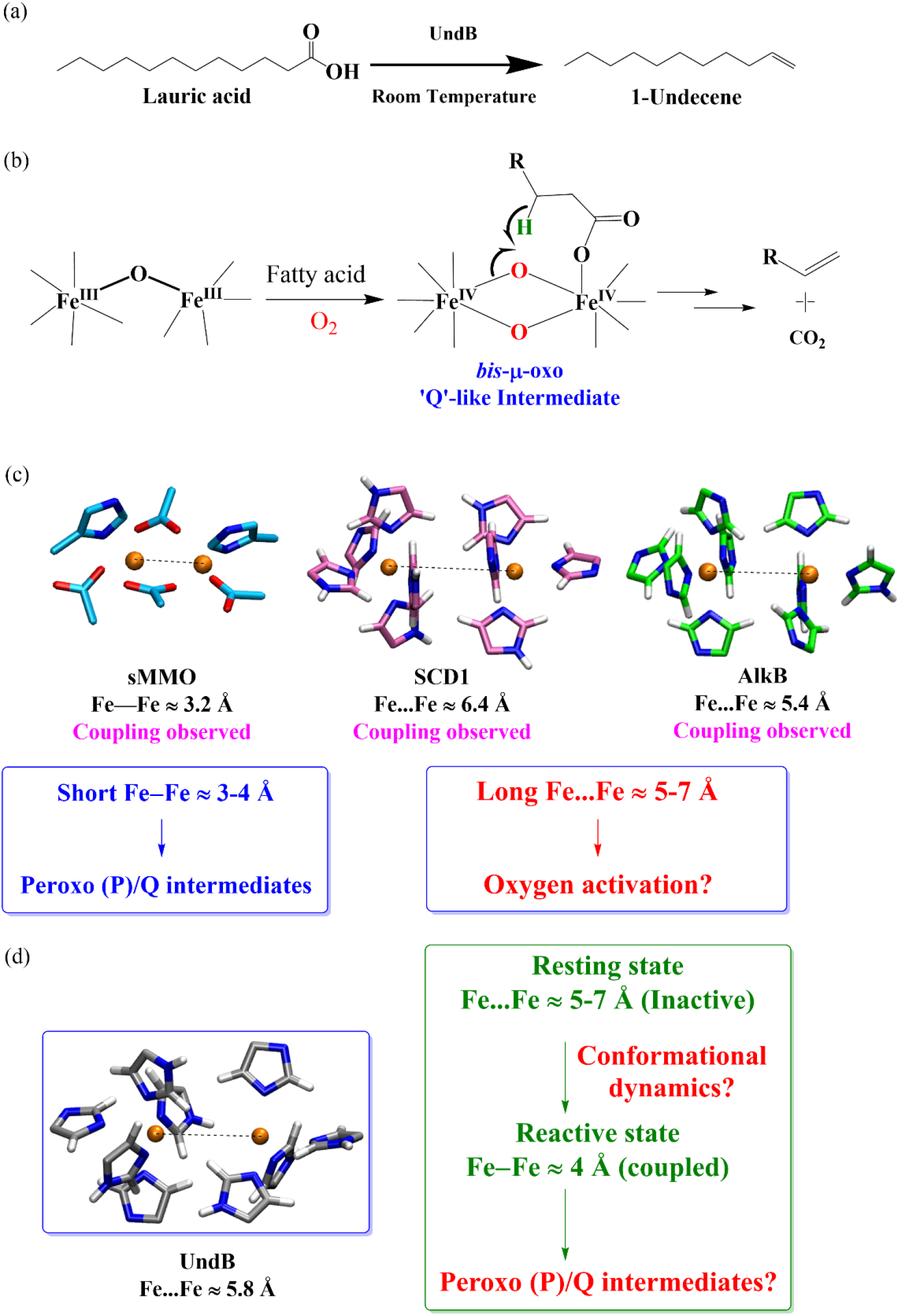
(a) The chemical reaction catalyzed by UndB. (b) Previously proposed mechanism for the conversion of fatty acids into terminal alkenes by UndB. (c) Structural–spectroscopic paradox: diiron active sites of soluble methane monooxygenase (sMMO), SCD 1 and AlkB, showing elongated Fe–Fe separations despite evidence of metal–metal coupling. (d) AlphaFold3-predicted active site of *Pc*UndB with an elongated Fe– Fe distance. schematic illustrating possible dynamic modulation between inactive (long-distance) and coupled (short-distance) states.

Since its discovery in 2015^9^, the membrane-bound fatty-acid decarboxylase UndB has remained mechanistically enigmatic, with only limited availability of structural and functional information^3^. Although UndB belongs to the fatty acid desaturase–like superfamily, which includes Stearoyl-CoA Desaturase 1 (SCD1) and Alkane monooxygenase AlkB and contains a characteristic nine-histidine motif that coordinates the diiron centre^10-12^ its chemistry diverges sharply from canonical desaturation. Kinetic studies have proposed a mechanism involving hydrogen atom abstraction (HAA) from the β-carbon of the substrate, followed by radical-mediated decarboxylation to generate terminal 1-alkenes (Fig 1b)^3,13^ The authors presented kinetic isotope effects, k_H_/k_D_=2.6, to establish the HAA as the rate-determining step (rds) in UndB compared to the Kolbe-type mechanism. In particular, UndB catalysis is accompanied by substantial hydrogen peroxide (H_2_O_2_) formation^14^, which is inconsistent with a tightly coupled high-valent oxidant such as the canonical diamond-core ‘Q’ intermediate commonly invoked for non-heme diiron enzymes. The previously proposed mechanism leaves several critical issues unresolved. In particular, it remains unclear how the two iron centers converge to form a ‘Q’-like intermediate within an active site constrained by nine conserved histidine residues.

These uncertainties are further amplified by recent structural and mechanistic studies of related desaturase-like diiron enzymes. Structures of SCD1^10^ and AlkB ^10,11,15,16^ reveal elongated Fe–Fe separations (≈ 5.4–6.8 Å), substantially longer than those typically observed in ferritin-like oxidase and oxygenase (FDO) enzymes such as soluble methane monooxygenase (sMMO) and ribonucleotide reductase (RNR), Δ9-Desaturase and Arylamine N-oxygenases AurF and CmII (Fig. 1c) where oxygen activation proceeds through well-defined peroxo (‘P’) and diamond-core (‘Q’) intermediates.

Shorter iron-iron distance is reported to be essential for O_2_ activation through the formation of ‘P’ and ‘Q’ intermediates in most of the non-heme diiron enzymes, making the understanding more confusing^5,17^. On the basis of these long Fe–Fe distances, alternative mechanisms have been proposed in which the two iron centres perform distinct roles during catalysis^4,16,18^. This mechanism not only lacks any experimental support, but also there are several experimental observations that it cannot explain. It becomes especially difficult to reconcile with consistent reports of coupled iron centres^1^, as is evident from Mössbauer and EPR spectroscopy in AlkB^2,15^ and SCD1^19^. While AlkB shows a strong presence of anti-ferromagnetic coupling (J < 80cm^-1^), SCD1 reveals a weak anti-ferromagnetic coupling (J ∼ 7 cm^-1^)^2,4^. The molecular origin of this coupling and how catalytically competent states emerge within these apparently elongated diiron centres without any bridging motifs, therefore, remains unresolved. Whether similar principles govern oxygen activation in UndB is currently unknown, especially in light of the anomalous H_2_O_2_ production and persistent spectroscopic evidence for iron–iron coupling in this enzyme family. These observations suggest that static structural models may not fully capture the conformational ensembles relevant for catalysis. In particular, dynamic fluctuations in the metal–metal distance could provide a mechanism for transiently establishing the electronic coupling required for oxygen activation (Fig. 1d). Testing this hypothesis requires a framework capable of resolving the interplay between protein conformational dynamics, electronic structure and catalytic energetics— an aspect that has remained largely unexplored for membrane-bound NHFe_2_ enzymes.

These inconsistencies raise fundamental questions^5,10^ about how membrane-bound diiron enzymes activate O_2_ and selectively cleave terminal C–H bonds, that have remained intractable to static structural and mechanistic approaches. To address this mechanistic problem, we investigated oxygen activation in the membrane-bound fatty-acid decarboxylase UndB employing an integrative multiscale computational framework combining large-scale atomistic molecular dynamics, quantum mechanics/molecular mechanics (QM/MM) molecular dynamics, free-energy simulations, high-level quantum chemical calculations, and AlphaFold3 modelling. Our simulations reveal that oxygen activation is dynamically regulated by fluctuations in the Fe–Fe distance, where transient contraction of the diiron centre establishes effective metal–metal coupling and enables formation of a catalytically competent peroxodiiron(III/III) intermediate despite elongated resting-state structures. Recently, this methodology was used to propose a new intermediate in SCD1 having a similar active site structure as UndB. Our multiscale simulations reproduce quantitative experimental activation barriers and explain major experimental findings. However, the pathway is considerably different from that of the other NHFe_2_ enzymes, in which the formation of a *cis*-1,2-peroxo intermediate was presumed unfeasible for SCD1 based on longer Fe…Fe distance and histidine-rich motifs^10^ and a recent study on AlkB^4,16^. These results reconcile structural and spectroscopic observations and suggest that dynamic metal–metal distance modulation constitutes a general mechanism governing oxygen activation in desaturase-like diiron enzymes. Our study also provides testable guidelines unexplored so far to enhance the catalytic efficiency of UndB through mutating certain residues. The outcome extends beyond elucidating the mechanism of UndB, establishing a general biophysical framework for extracting reliable mechanistic insights from membrane-bound enzymes and reconciling longstanding discrepancies between structural and solution-phase behavior.

## Results and Discussion

### Active-site organization of the enzyme–substrate–O_2_ (ESO_2_) complex

Structural and spectroscopic studies of non-heme diiron enzymes have revealed an apparent mechanistic paradox: whereas spectroscopic measurements indicate coupling between the two iron centres during catalysis, most available structures display elongated Fe–Fe separations that appear incompatible with such coupling. To examine how this discrepancy arises in the membrane-bound fatty-acid decarboxylase UndB, we first established a structural model of the enzyme–substrate– O_2_ (ESO_2_) complex. Given the limited experimental structural information available for UndB, we employed AlphaFold3 (AF3) to generate three-dimensional structures (3D) incorporating the diiron cofactors^20^. Alphafold3 has previously been successfully applied to predict the 3D structures of enzymes notably membrane enzyme such as AlkB^4^. In this study, the structures were predicted using homologous sequences from *Pseudomonas canadensis* (PcUndB) and *Pseudomonas mendocina* (PmUndB) which share 85% sequence similarity. Both models yielded identical architectures having Fe…Fe distance of 5.87 Å (Fig. 1d). Since both UndB structures have led to similar results, which establishes the robustness of the study, the following results are discussed with respect to PcUndB unless otherwise stated. QM/MM free energy simulations of the substrate-unbound UndB gives the most stable Fe…Fe distance to be 5.7Å as discussed later (Methods, section 1.4, SI). The AF3-predicted UndB structure is in good agreement with biochemical characterization that the active site of UndB consists of two non-heme iron ions coordinated entirely by nine histidine residues much like SCD1 and AlkB^3^ (Fig. 1c) as well as confirming it is an O_2_-dependent enzyme. The nature of the substrate-binding channel is further revealed through a subsequent enzyme engineering experiment thus confirming two important sites: the active site and substrate channel nature in UndB^21^.

Extensive atomistic MD simulations (1.8 μs cumulative sampling) of multiple UndB replicas, revealed two dominant Fe…Fe distance populations at 5.10 Å and 6.42 Å (Fig S7, SI) in the substrate-O_2_-bound UndB. The distance between the iron centers falls in line with that of SCD1^22^ and AlkB which was reported to be 6.4-6.8 Å and 5.4-6.1 Å. These observations indicate that the active site architecture is globally conserved and remains structurally stable, as reflected by the comparable inter-iron distances observed across the family. However, such separations are difficult to reconcile with the consistent metal–metal coupling inferred from spectroscopic studies, suggesting that the MD-sampled resting-state ensemble does not correspond to the catalytically competent configuration. Moreover, the binding orientation of the substrate, lauric acid, closely matches with the experimentally characterized substrate pose in UndA^8^. However, a reactive conformation of UndB was not obtained despite extensive conformational sampling, as evidenced by the persistently large separation between the bound O_2_ molecule and the substrate’s β-hydrogens (β-H) atoms (Fig. S8, SI).

Consistently, QM/MM molecular dynamics simulations of UndB with lauric acid and O_2_ bound to either Fe centres mostly fails to produce any reactive geometry^5^ (Fig. 2a,b) in the relevant high spin S = 4, 5 states that was reported for non-heme diiron enzymes and was also considered in recent studies on SCD1 and AlkB having similar active site structure^23,24^ (Fig. S9, SI). Throughout the 60 ps QM/MM simulations, the β-H remains distal from the bound oxygen atoms (Fig. 2b and Fig. S9, SI), whereas molecular O_2_ preferentially binds to or transiently hovers over the substrate-bound Fe_B_ centre, revealing its role as the initial activation site. Substrate binding other than coordinating to a Fe centre gives rise to steric clashes with the active site residues. A similar non-reactive conformation has been observed in the lauric acid-bound UndA crystal structure, raising the question of how the reaction is subsequently initiated^8^.

**Fig. 2.**
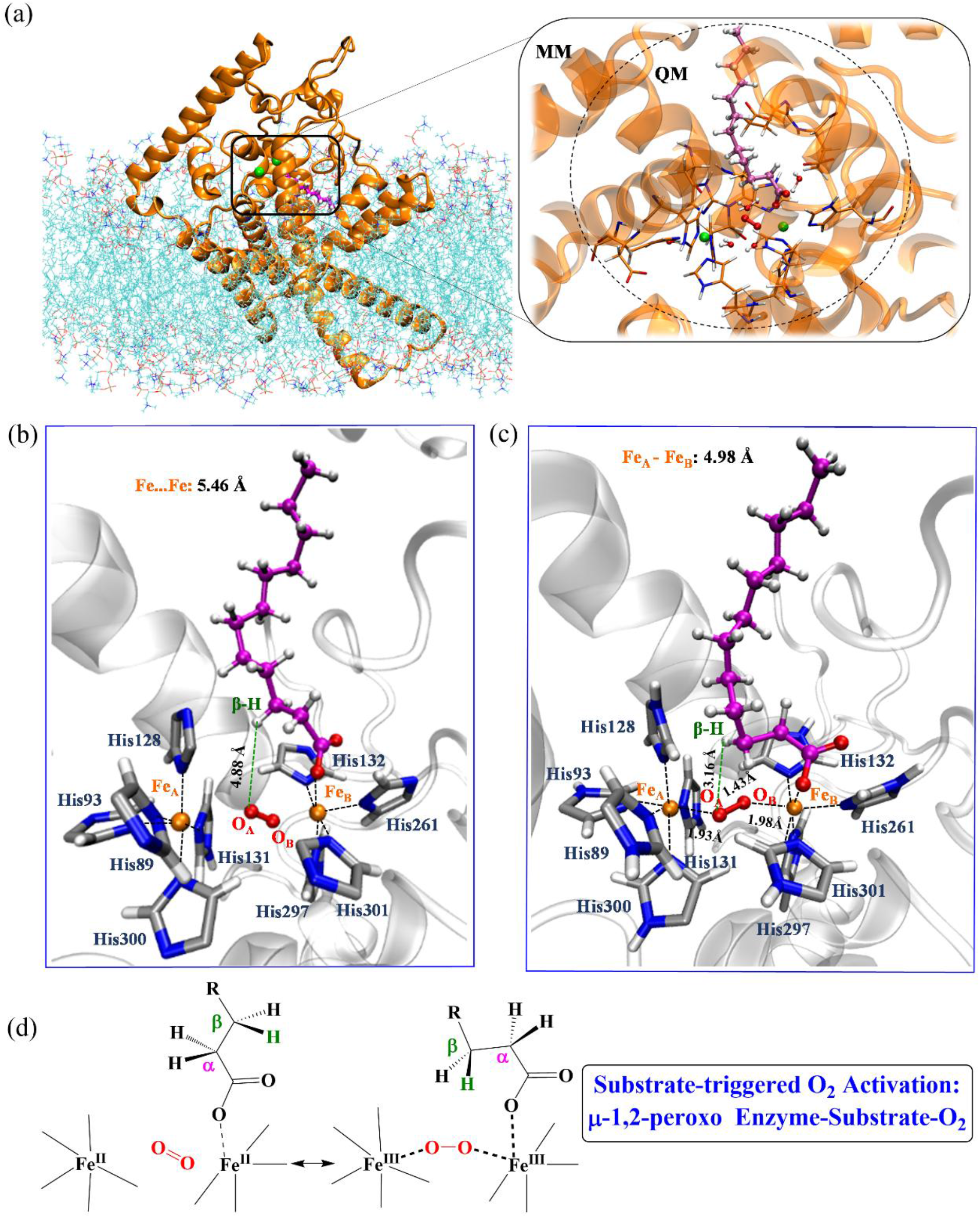
(a) UndB embedded in POPC lipid membrane; (inset, right) setup of the QM/MM simulations and a closeup of the 312 atoms included in the QM region of QM/MM MD simulations. (b) the active site region of the enzyme-substrate-O_2_ complex from QM/MM MD simulations. (c) Peroxodiiron(III/III) intermediate as obtained from the specialized QM/MM MD simulations. (d) Schematic representation of the µ-1,2-peroxo species formation from substrate-triggered O_2_ activation.

The elongated Fe–Fe separation has emerged as a structural signature of histidine-rich non-heme diiron enzymes, in contrast to the shorter distances observed in ferritin-like diiron oxidase and oxygenase families^5,17^. Such an expanded metal–metal separation suggests a fundamentally different mode of O_2_ activation during catalysis^6,10^. How molecular oxygen is activated by two spatially separated iron centres therefore remains unresolved, raising the possibility that transient structural rearrangements dynamically establish the metal–metal coupling required for catalysis.

### Substrate-triggered oxygen activation forms a peroxodiiron(III/III) intermediate

Considering the inactive conformation obtained from high-spin QM/MM simulations, we employed a variant of steered molecular dynamics (SMD) simulations called conditional SMD (c-SMD-QM/MM MD) to induce the possible reaction mechanisms. An intermediate structure was selected from the SMD trajectories. This method efficiently explores the reaction pathway of UndB in the absence of any prior knowledge of the active state^25^. The c-SMD-QM/MM MD simulations was followed by 100 ps of QM/MM MD run. The QM region consists of 312 atoms that involves the diiron center (Fe^2+^), active site histidines (Asp 90, Glu142, Glu 143, (His89, His93, His128, His131, His132, His261, His297, His300, and His301)^9^, the lauric acid substrate, and water molecules within 2.5 Å of the active site (Fig. 2a). The unbiased QM/MM MD followed after the near-attack conformation obtained from the above SMD run showed a successful abstraction of β-hydrogen by the activated oxygen species and release of carbon dioxide within 15 ps, and the resulting conformation was found to be stable throughout the 100 ps long simulation in high spin S = 4 state (see Section 3.0, SI). Considering the limitations of semi-empirical QM method, the QM/MM MD simulations were performed with high-spin state S = 4 with all unpaired electrons aligned in the same direction that allows us to investigate the same electronic configuration of all the structures as with DFT in view of longer Fe-Fe separations. This formalism has been employed in a recent study on SCD1 having similar active site structure to a capture a new intermediate^23^. It may be noted that the target of the QM/MM MD simulations is to explore all dynamically accessible intermediates of the ESO_2_ complex in the enzyme environment considering conflicting experimental observations that could not be captured by classical simulation approaches. However, the chemical viability of the corresponding electronic structures was studied using independent high-level quantum chemical calculations and strong correlation with experimental observations. In this, we constructed a cluster model of UndB consisting 158 atoms from the appropriate QM/MM MD snapshot that includes two iron ions, nine coordinating histidine residues and a lauric acid substrate for investigation using a high-level quantum chemical method (Fig. S2, SI).

QM/MM MD simulations generated one reactive conformation of UndB: a peroxide bridging species, similar to the ‘P’ intermediate reported for sMMO, not its famous successor ‘Q’^5^. It also shows product (1-alkene) formation along with the release of carbon dioxide (CO_2_). The formation of the peroxodiiron(III/III) intermediate is supported by *ab initio* quantum chemical density functional theory (DFT) calculations (see Section 3.0, SI). That calculations revealed any attempt to optimize a ‘Q’-like intermediate results in a bis-Fe-oxo intermediate, which however showed significant high barrier for β-H abstraction.

All other O_2_ binding to either of the Fe centres converge eventually to the peroxodiferric ESO_2_ complex in this the multiscale approach. QM/MM MD simulations reveal O_2_ binds preferably to the Fe_B_ ion coordinated with the substrate, LA, which is also supported by quantum chemical DFT calculations with anti-ferromagnetically coupled Fe centers. Without the LA binding, O_2_ does not bind to either of the Fe centres and is not activated for chemical reaction as revealed by longer Fe– O distance (2.30 Å) (Table S1, SI). It implies that this is a case of substrate-triggered O_2_ activation, as observed in UndA^8^, ADO^26^ and in other enzymes^5^. The Fe_B_ centre bound to the substrate transfers the first electron to the molecular O_2_ for its activation, supporting a substrate-triggered activation mechanism (Table S2, SI). It may further be noted that a superoxo-like ESO_2_ complex with O_2_ bound to either of the Fe centre is only observed in the high spin state at longer Fe-Fe distance, albeit with O_2_ less activated than the peroxide intermediate (Section 3.1, SI). Abstraction of a hydrogen atom from a strong C–H bond of lauric acid (BDE > 95.0 kcal/mol) by a metal superoxo species appears less likely^8,27,28^, which is supported by quantum chemical calculations discussed in section 3.0, SI. Addition of superoxide dismutase (SOD) in the reaction media does not affect the UndB catalysis, while catalase addition profoundly affects UndB catalysis, indicating production of peroxide, not superoxide species in the media^14^. The QM calculations show that O_2_ is activated in the peroxo-bridged state in UndB, not in any other form as suggested for SCD1 and AlkB^18,24^. To obtain the activation barriers of the HAA step, we performed QM/MM well-tempered metadynamics of the major catalytic steps.

The formation of a μ-peroxodiiron(III/III) intermediate is particularly striking in UndB, given its unusually long Fe…Fe separation and the absence of canonical bridging ligands such as carboxylate or oxo/hydroxo groups. Shorter Fe…Fe distance (3.0 - 4.2Å) and presence of bridging ligands, as in sMMO, RNR and other FDOs, have been a structural hallmark for peroxodiiron(III/III) intermediate formation^5,17^. The peroxo-like intermediate obtained from QM/MM MD and QM calculations are shown to have Fe-O distances (Fe_A_-O_A_ = 1.93 Å and Fe_B_-O_B_ = 1.98 from QM/MM MD (Fig. 2c) and 1.96 Å and 1.98 Å respectively at the B3LYP-D3/def2-SVP level) relatively longer than experimentally observed Fe–O bond lengths (≈ 1.88 Å) in diferricperoxo intermediate^5^. It points towards a weakly bound or quasi-stable peroxodiferric complex. This intermediate could possibly leak out in the solution media to produce hydrogen peroxide (H_2_O_2_). This is supported by *in situ* production of large amounts of H_2_O_2_ during UndB catalysis^14^. Detection of H_2_O_2_ has been an important indicator for peroxide intermediate formation in NHFe_2_ enzymes^5,29^. Characterization of such peroxo species in SznF, HrmI and Δ9-Desaturase endowed with longer Fe…Fe distance (> 4.0Å) lends strong credence to it (Fig. 4c). This expands the implications of the present work beyond the desaturase-like family to other enzymes class. This kind of peroxo intermediate without any oxo-bridge is reported for heme-oxygenase-like diiron oxidase and oxygenases (HDO) metalloenzyme SznF^30-32^. Thus, for the first time our study unravels the possible origin of large amount of H_2_O_2_ production during UndB catalysis and thereby could suggest pathways to mitigate it for effective catalysis.

### Dynamic Fe–Fe distance modulation enables formation of the reactive oxidant

QM/MM MD simulations reveal that the μ-peroxodiiron(III/III) intermediate is itself incapable of abstracting the β-hydrogen from lauric acid. Instead, elongation of the O–O bond—facilitated by a nearby water molecule acting as a proton donor—promotes O–O bond cleavage and leads to the formation of a high-valent iron-oxo (Fe(IV)=O) species. This reactive iron–oxo unit, generated via a proton- and electron-assisted transformation of the peroxo intermediate into a hydroperoxo-derived state, is competent for β-hydrogen abstraction from the substrate. This was followed by QM/MM free energy simulation employing well-tempered metadynamics to determine the activation barriers of the major catalytic steps of UndB in high spin state (with S = 4.5) taking into account the additional proton and electron. The O–O The bond-breaking step shows an activation barrier of 10.7 kcal/mol^33^ (Fig. 3a,b), and the activation barrier for β-H abstraction by the iron-oxo species is 14.1 kcal/mol, which is calculated to be 15.0 kcal/mol from QM calculations. This is in good agreement with the experimentally obtained barrier of 17.9 kcal/mol (from experimental k_cat_), considering the usage of semi-empirical methods^23^. Our QM/MM free energy calculations also show the β-H abstraction step as the rate-determining step (rds), in agreement with experimental kinetic data. The long iron-iron distance in membrane-bound enzyme UndB implies weak exchange coupling; the high-spin QM/MM free-energy surface captures the dynamical formation and reactivity of the peroxodiiron intermediate, as independently supported by the substantial H_2_O_2_ formation observed experimentally as well as high-level QM calculations. Essentiality of an extra electron and a proton for UndB catalysis is supported by the consumption of four reducing equivalents (NADPH). More support is drawn from a recent study of Arylamine Oxygenases and related binuclear NHFe_2_ as well as soluble Δ9-Desaturase enzymes, where the supply of electrons and protons was shown to be indispensable for their catalysis^33,34^. The viability of the electronic structure of the key intermediates was checked by cluster model QM calculations. Major catalytic steps involve the formation of a (hydro)peroxo intermediate that generates the active iron-oxo species, which then performs the rate-determining β-H abstraction followed by decarboxylation to generate the product (see Section 3.0, SI). Consistency between the QM/MM free-energy profiles and independent electronic structure calculations supports a peroxodiiron-mediated C–H activation pathway.

**Fig. 3.**
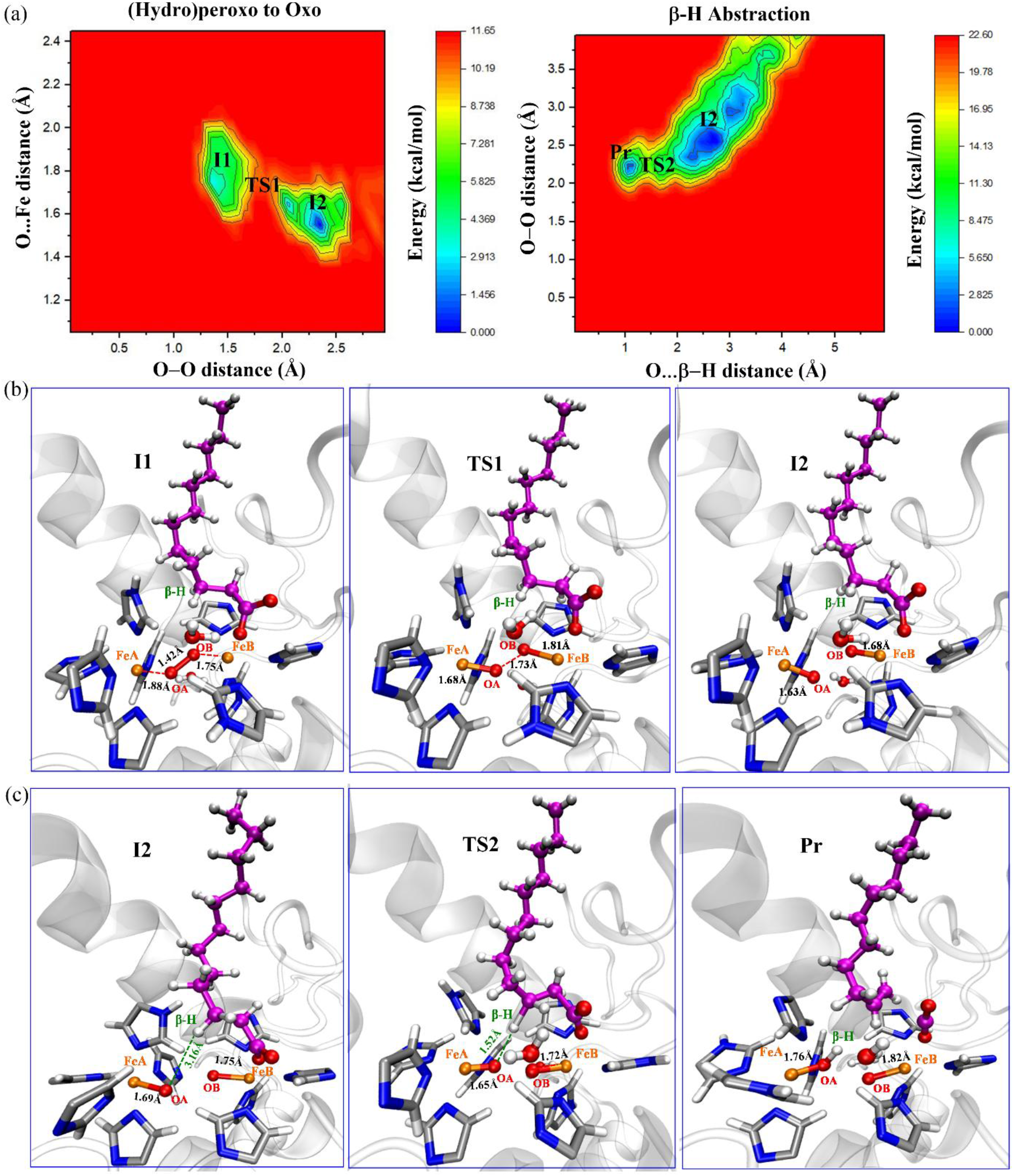
QM/MM free energy landscape of the (a) O–O bond breaking step and the rate determining β-H abstraction step in UndB at PM6/CHARMM36 level. (b) Possible intermediates and transition states obtained during the free energy simulations for (b) O–O bond breaking step and (c) the β-H abstraction step from the substrate, lauric acid in UndB.

Interestingly, we observed a striking feature in the Fe..Fe distance. During the breaking of the (hydro)peroxo O–O bond, the Fe…Fe distances samples a relatively wide range (4.6Å-6.6Å) of distances (Fig. 4a). It reveals that the two iron cofactors are capable of contracting in the presence of O_2_ and substrate. Attempt to decrease the iron-iron distance without the substrate and co-substrate below 5.0 Å leads to a barrier of 13.0 kcal/mol and above (Fig. S11, SI), a result also supported by a combined computational and experimental study on AlkB^4^. Multiscale dynamical simulations can capture such a peroxodiiron species, which is not accessible in classical simulation methods. Moreover, efforts to obtain an oxo or hydroxo-bridged structure didn’t succeed even after several attempts giving more support for a peroxo-bridged P-like species rather a Q-like species, similar to QM calculations (Section 3.0, SI). The dynamic nature of the iron cofactors also draws support from multiple iron-iron distances reported from different resolved structures of AlkB and SCD1. Generally shorter interatomic distance between the iron centers is essential for the formation of the peroxodiiron(III/III) intermediate, reported in several studies^5^. In this context, our study illustrates that iron-iron coupling leading to catalytic competence can emerge despite the absence of a canonical bridging motif. It thus appears that nature uses this elegant strategy to reduce the O–O bond breaking barrier in order to perform such an energetic C–H bond breaking reaction. This possibly reveals why and how peroxodiiron(III/III) species is reported in spite of long iron-iron distances, e.g., in HDO enzyme SznF^31,32^.

**Fig. 4.**
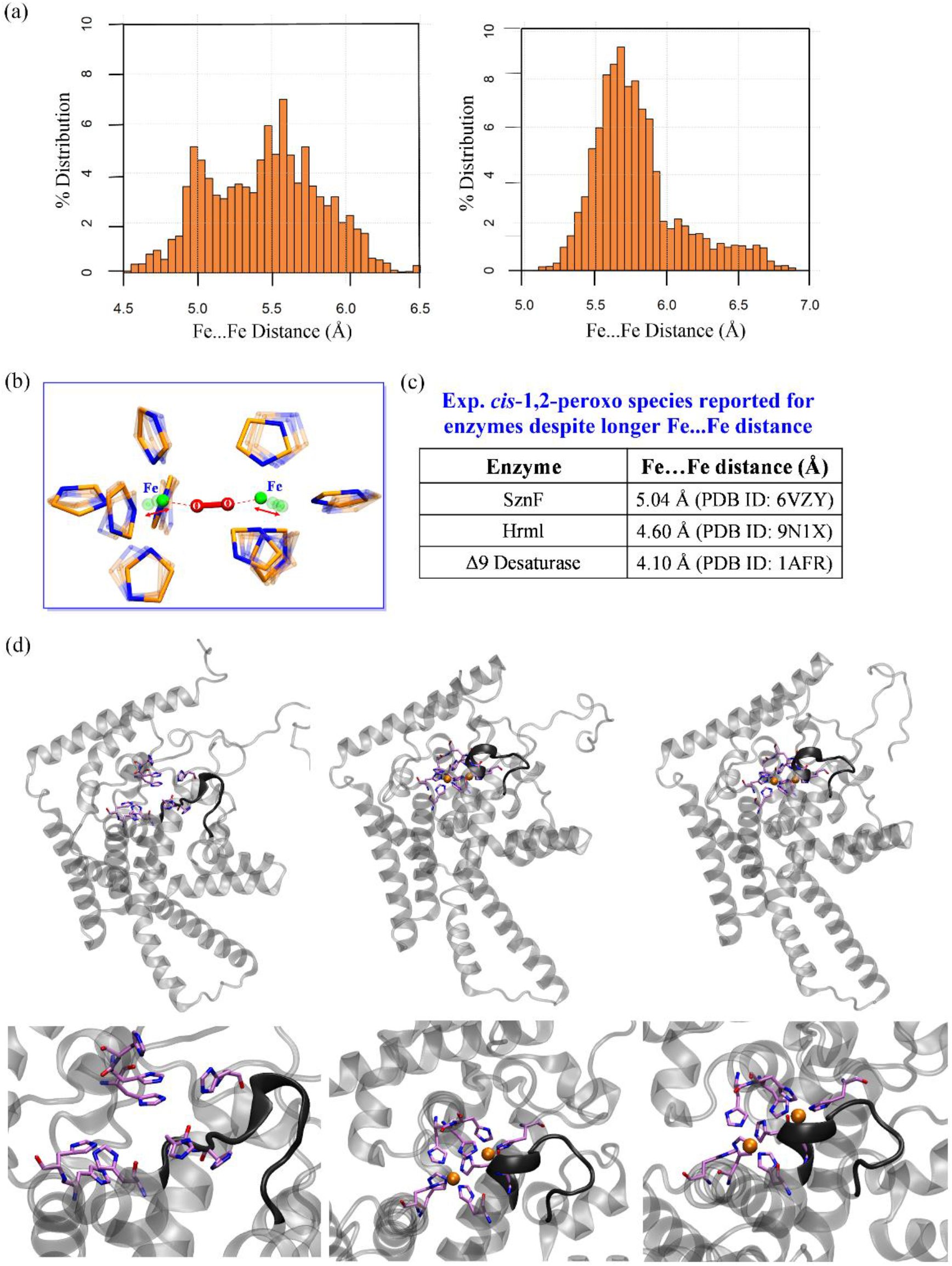
(a) Histogram population analysis of Fe…Fe distances during QM/MM metadynamics simulation trajectories of the PcUndB. (b) Dynamic Fe cofactors from QM/MM free energy simulations. (c) Enzymes with experimental report of peroxodirron(III/III) intermediate formation despite longer Fe…Fe distance.(d) Structural comparison of the (a) apo-, Fe(II)_2_- and substrate-bound UndB from large-scale classical simulations. Winding-unwinding of the iron-binding helix 5 (residues 132-141 colored black) shown for the above three structures respectively.

This work also addresses a long-standing discrepancy between the structures of this family of enzymes and conflicting spectroscopic observations, where the two iron centers are reported to be coupled through some unknown mechanism. It is this anti-ferromagnetic coupling of the two iron centers through a dioxygen-derived peroxo bridge that most possibly explains the coupled diiron cofactors with an anti-ferromagnetic (AF) coupling constant of J ≈ 15 cm^-1^ obtained from high-level QM calculations. The value of J (15 cm^-1^) falls between the J values of AlkB (J ≈ 80 cm^-1^: strong AF) and SCD1 (J ≈ 7 cm^-1:^ weak AF), which is also in harmony with the respective iron-iron distances in these enzymes. The coupling constant matches those values for dioxygen-derived bridges as reported by Solomon and coworkers^33^. The quasi-stable nature or fleeting existence of the peroxodiiron(III/III) species is probably responsible for its escape detection by any method. Considering the anti-ferromagnetic coupling constant value of UndB obtained from QM calculations, we also posit that such dynamic iron cofactors exist in AlkB and SCD1, which result in coupling of the two iron centers in the absence of any bridging ligand and consequently exhibit characteristic EPR and Mossbäuer spectra^4,16^. Our study thus addresses a long-standing mystery that existed between the structures of AlkB & SCD1 family of enzymes and their solution behavior.

Efforts were also made to understand the underlying rationale behind shortening of the Fe…Fe distance, as this is possibly a structural hallmark for most NHFe_2_ enzymes. From classical simulations of apo-, Fe(II)_2_-UndB and substrate-bound UndB, we noticed an unstructured/loop region for residues 132-141 (Fig. 4d). This disordered region becomes helical (helix 5:H5) on binding with the Fe cofactor, which remains more or less unchanged in the substrate-bound UndB. The winding-unwinding nature of H5 makes the substrate-bound Fe_B_ center flexible enough for shortening of the Fe…Fe distance in substrate-bound UndB. The disorder-order transition in the iron coordination site is emerging as a unifying characteristic similar to programmed instability observed in SCD1 and HDOs like UndA^4^, SznF^32^. Based on this, we conjecture that site-directed mutations like G136P, T137V, S135K and T139E may provide a more regulated diiron coordination site, thereby shortening the iron-iron distances enough to form a stable peroxo species for effective UndB catalysis. Such an engineering strategy may also help to decrease the reducing equivalent required, a previously coveted feat^21^. To get more insight into the conformational transition between the active and inactive states of UndB, we turn our focus next on this matter.

### Conformational transition to the catalytically competent state

Classical metadynamics simulations reveal that the conformational transition to the active state from the inactive state has a free energy barrier of 6.0 kcal/mol (Fig. 5a,b), which is 4.1 kcal/mol obtained from QM calculations. The substrate tunnel shape does not change during the active state transition, as seen in the free energy landscape constructed with Arg124-Glu143 distance. Importantly, the active conformation orients the β-H of lauric acid towards the activated oxygen atom of molecular O_2_; at the same time, the α-Hs are held back away by the Glu143 residue from the activated oxygen species. QM/MM MD and QM simulations have shown LA has a strong binding preference with the tetra-histidine coordinated Fe_B_ center and the substrate remains attached to it until the decarboxylation step (see Section 3, SI). These support the biochemical results that have clearly revealed a regiospecific product, 1-undecene, resulting only from β-H abstraction from the substrate lauric acid^3^.

**Fig. 5.**
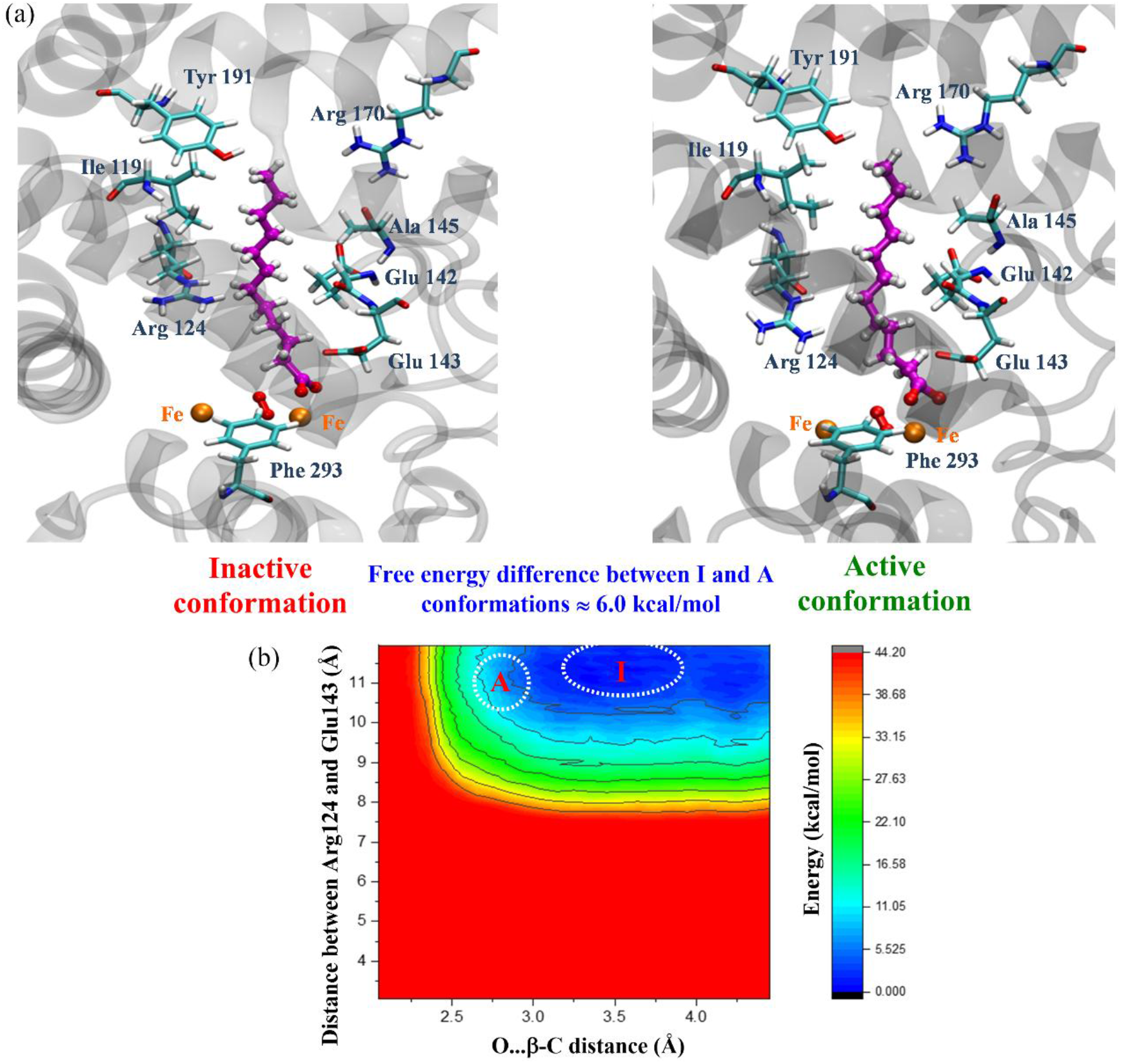
(a) Snapshots of the inactive and active UndB conformations obtained from QM/MM MD simulations at the PM6/CHARMM level. (b) Free energy surface for the conformational transition from the inactive to the active state from metadynamics.

Among all other substrate channel residues, Arg124 and Glu143 were found to be very important for stabilizing the active conformation through primarily electrostatic interaction^21^ (Fig. S15, SI). Apart from guiding the incoming substrate LA to the active site diiron centers, the two residues envelope the substrate, LA, from the opposite side to stabilize the active conformation. These interactions help in keeping the active site residues in a preorganized state hence facilitating catalysis in UndB, similar to AlkB^11^. Drastic reduction of the enzymatic activity of the R121F mutant equivalent to R124^21^ provides strong support for it, as a bulky phenylalanine residue would hinder the necessary conformational transition to the active state.

In that study, a domain swap engineering strategy was reported for UndB, supported by classical MD simulation studies for the conversion of long-chain fatty acids, which corroborate the bulky nature of the substrate channel^21^. Moreover, alanine scanning E139A and R121A mutants equivalent to E143 and R124 also show significant reduction in activity^35^. This reinforces the requirement of a preorganized active site^36^ which is achieved by stabilizing the active conformation for effective catalysis. Alanine residue is smaller to maintain such necessary substrate channel shape. Based on this insight, it is prudent to make R124K point mutation for proper anchoring of the substrate and assistance in maintaining the correct preorganization active site in order to enhance the catalytic efficiency of UndB to meet industrial standards^37^. Other substrate-binding residues, such as Ile119, Glu142, Ala145, Tyr191, and Phe293, are also important for accommodating the conformation flexibility that leads to successful catalysis. Among other nonheme diiron enzymes, the soluble UndA enzyme employs substrate tunnel residues of similar nature to bind lauric acid^11^. The flexible nature of the incipient substrate radical directly bound to the Fe_B_ center would be helpful to thwart any OH rebound step.

Similar arguments were put forward in the UndA case for explaining the β-H abstraction regiospecificity^38^. Indeed, such active conformation of UndB draws support from the β-hydroxy dodecanoic acid (BHDA) bound UndA crystal structure, where the β-hydroxyl oxygen is positioned only 2.45Å away from the distal oxygen atom^7^ without any noticeable change of the protein scaffold, as is also observed in UndB with the same substrate LA. This is in contrast to the OleT enzyme, where the substrate is precisely held in position for an effective OH rebound reaction^39^. In this context, the abstraction of another H (either from α or γ position) for the formation of an unsaturated product seems not feasible due to the respective hydrogens not being within the reacting distance of the reactive oxygen species. Thus, direct binding of flexible C-12 fatty lauric acid to the Fe_B_ centre is most likely the main reason for maintaining regiospecificity, frustrating OH rebound and preventing another H abstraction in UndB, thereby revealing the importance of substrate dynamics in effective UndB catalysis^40^.

## Conclusions

Our study demonstrates that oxygen activation in desaturase-like non-heme diiron enzymes is governed by transient conformational dynamics rather than static active-site geometry, as revealed through a detailed investigation of UndB, thereby providing a resolution to longstanding discrepancies between structural and spectroscopic observations. We establish a dynamically gated mechanism in which transient Fe–Fe contraction enables effective metal–metal coupling and formation of a quasi-stable μ-1,2-peroxodiiron(III/III) intermediate, thereby facilitating O_2_ activation despite the absence of canonical bridging motifs. This framework reconciles kinetic, spectroscopic and energetic observations, including the formation of H_2_O_2_ during catalysis. More broadly, our results suggest that flexible diiron cofactors may represent a general mechanistic feature of histidine-rich diiron enzymes, bridging mechanistic understanding across systems such as UndB, AlkB and SCD1, and providing conceptual links to related diiron systems such as soluble methane monooxygenase. These findings provide a general biophysical framework for linking protein dynamics to catalytic function in membrane-bound metalloenzymes and open new avenues for rationally tuning iron–iron distances through targeted mutations and engineering catalytic efficiency in biofuel-relevant enzymes. The dynamically coupled diiron mechanism uncovered here provides a unifying framework for understanding how nature harnesses O_2_ for demanding chemical transformations and defines design principles for controlling oxygen activation in complex metalloenzymes.

## Methods

Large-scale atomistic molecular dynamics (MD) simulations establish the structural stability and conformational flexibility of the active site, providing a realistic description of the membrane-bound enzymatic environment. Semi-empirical QM/MM MD simulations establish the dynamical accessibility of key intermediates within the enzymatic environment, which are difficult to capture using static methods. Free-energy calculations identify the rate-limiting chemical step and confirm consistency with experimentally measured activation barriers, while independent cluster model QM calculations demonstrate the chemical viability of the corresponding electronic structures. Although absolute energetics depend on the chosen Hamiltonian, all approaches converge on a consistent mechanistic framework, which is further supported by experimental observations. Together, these results reveal a peroxodiiron(III/III)-mediated O_2_ activation pathway in UndB that contrasts with the previously proposed canonical mechanism.

Considering the remarkable success in predicting correct 3D structures of proteins from their amino acid sequences, we have used Alphafold3 to generate an initial structure of UndB from its amino acid sequences. The initial structure for model preparation was generated using AlphaFold3^20^ using the (UniProt ID: A4Y0K1 and GenBank: UKZ44198.1), with Fe ions incorporated into the active site through the AlphaFold3 method. From the sequences generated by AlphaFold3, the structure with the highest confidence was selected for system preparation. The substrate lauric acid was docked and incorporated into the protein using the ligand binding pose from UndA^8^ [PDB: 6P5Q]. An oxygen molecule (O_2_) was also inserted into the active site region to prepare an enzyme-substrate-O_2_ complex for investigation. A total of 1.8 µs of all-atom simulations were performed using a timestep of 1 fs. Trajectory analysis was performed using the VMD^41^ program. Molecular dynamics (MD) simulations were performed using the GPU-accelerated version of the NAMD^42^ package. The simulations were run for 300 ns for each replica of UndB.

The QM/MM interface of NAMD^43^ was used to select the QM region and initiate the QM/MM simulation using the QwikMD interface^44^ provided by VMD. The PM6-D3H4^45^ method was employed for the QM region using the NAMD-MOPAC interface^43^, while the MM subsystem was treated with the CHARMM36 force field^46^. A total of 460 ps of QM/MM MD simulations were ran. We created the cluster models^18,47^ for quantum chemical calculations from appropriate snapshots of the QM/MM MD simulation trajectory. All quantum chemical (QM) density functional theory (DFT) calculations were carried out using the ORCA program^48,49^

A variation of steered molecular dynamics (SMD) simulations called conditional SMD (c-SMD)^43^ was performed to induce the possible reaction mechanisms, and an intermediate structure was selected from the SMD trajectories. A moving harmonic restraint was applied along the reaction coordinate defined as the distance between the activated oxygen and the β-hydrogen of the lauric acid (C12) substrate. Upon reaching the cutoff distance, the restraint was released and the simulation was continued as unbiased QM/MM molecular dynamics. The QM region for all the QM/MM MD simulations consists of the diiron center (Fe^2+^), active site histidines (His89, His93, His128, His131, His132, His261, His297, His300, and His301), the lauric acid substrate, and water molecules within 2.5Å of the active site. The system was subjected to initial MM minimization using a conjugate gradient algorithm, followed by sequential QM/MM minimizations and equilibration through the QwikMD interface. The VMD-QwikMD interface was used for system setup^44^. The total charge and spin multiplicity were set to +1 and 9, 11 respectively corresponding to total spin states, S = 4 and 5. All QM/MM MD, and free-energy simulations were performed on a high-spin diiron surface. The equilibrated intermediate structure from c-SMD was used as an initial structure for QM/MM well-tempered metadynamics (WT-MetaD)^50,51^ simulations to enhance sampling along the collective variable space. All other simulation parameters were identical to those used in classical QM/MM molecular dynamics. The initial structure for classical metadynamic^52,53^ simulation was taken from the lauric acid-bound UndB after equilibration. An extended method section is available in the supporting Information. This includes details of the atomistic molecular dynamics simulations, metadynamics, QM/MM MD simulations, QM/MM well-tempered metadynamics and quantum chemical calculations.

## ASSOCIATED CONTENT

### Supporting Information

Extended Methods and Cartesian coordinates of calculated structures.

## AUTHOR INFORMATION

### Author Contributions

S. R. designed research, S. N. and A. C. performed research, S.R., S. N. and A. C. analyzed data, S.R., S. N. and A. C. wrote the paper.

## ACKNOWLEDGMENT

S. R. acknowledges support from Ramalingaswami Re-entry Fellowship (BT/RLF/Re-entry/39/2017), Department of Biotechnology, India. S. N. acknowledges doctoral fellowship from DBT, Govt. of India and A.C. thanks SINP for her doctoral fellowship. We thank Dr. Debasis Das, IISc, Bangalore for giving the idea of UndB enzyme along with helpful discussions. The financial support from DAE-Saha Institute of Nuclear Physics, Kolkata, (RSI-4002) is duly acknowledged. We thank the SINP computing facility for the computational resources.

## Data Availability

All data are available in the main text and SI Appendix.

## Competing Interest Statement

The authors declare no competing interests.

## Notes

### Competing Interest Statement

The authors have declared no competing interest.

### Summary of Updates

This version revises the mechanistic framework by highlighting dynamic Fe-Fe distance modulation in oxygen activation, with improved writing, analysis and figures.

