## Supporting Information for "Dynamic Metal–Metal Distance Modulation Controls Oxygen Activation in Non-heme Diiron Enzymes"

**This word file includes:**

Methods
Figs. S1 to S16
Tables S1 to S4
Cartesian coordinates of the optimized structures
SI references

### 1.0 Methods

#### 1.1 Molecular Dynamics (MD) Simulations

Due to the absence of experimentally resolved structural data for UndB, the initial structural model employed in this study is generated using AlphaFold3[1], with UniProt accession A4Y0K1 and GenBank entry UKZ44198.1 provided as the input sequences. The Fe ions are incorporated into the putative active site during model generation. Among the predicted models, the structure with the highest overall confidence was selected for subsequent system preparation. The model confidence, as assessed by predicted local distance difference test (pLDDT) scores, indicated very high reliability ($\geq$90) for the core transmembrane helices and structured domains,70–90 for surface helices and amphipathic segments, and <70 (low confidence) for flexible loops and terminal regions. The final models yield an Fe…Fe separation of 5.87 Å for both UndB structures. The structural integrity of AF3-predicted UndB has been checked (Section 2, SI). Two slightly different sequences were used to ensure that the active site was conserved in both cases.

The substrate lauric acid was docked and incorporated into the protein using the ligand binding pose from UndA[2] [PDB: 6P5Q]. An oxygen molecule ($O_2$) was also inserted into the active site region and was held there using a minimal patch; patches were also used to keep the ligand in the proper orientation. The protein–substrate–Fe incorporated system was then inserted into a lipid bilayer made of 1-palmitoyl-2-oleoyl-sn-glycero-3-phosphocholine (POPC), most commonly seen ER membrane lipid, with the initial orientation of the protein taken from the OPM server[3]. The membrane was built using the Membrane Builder plugin in VMD[4]. The system was then solvated with TIP3P[5] water box, and after solvation, it was neutralized with 0.15 M NaCl in a periodic box of 110 × 110 × 120 Å. The topology parameters for the substrates were generated using the ParaTool plugin of VMD, and the force field was generated using the CHARMM General Force Field (CGenFF)[6]; for the rest of the system, the topology and parameters were generated using the CHARMM36 force field[6].

The prepared system was first subjected to a lipid annealing step (allowing the lipid molecules to surround the protein as they would in a biological system), where the entire system (including the phosphate headgroups of the lipids) except the lipid bilayer was constrained with a force constant of 1 kcal/mol·Å$^2$ for 0.5 ns, ensuring no water penetration into the membrane. Following lipid annealing, a force constraint of 7 kcal/mol·Å$^2$ was applied to all atoms within 5 Å of the Fe atom. The system was then subjected to cycles of minimization (500,000 steps) followed by equilibration (200,000 steps) using a constraint-scaling approach, with the constraints gradually reduced as the simulation proceeded. The total equilibration time was 1.2 ns. After equilibration, the active site restraint was scaled up to 25 kcal/mol·Å$^2$, followed by a production run of 300ns. The same protocol was used for both *Pc*UndB and *Pm*UndB structures, incorporating lauric acid, two Fe-bound structure, two replicas of the *Pm*UndB structure, two replicas of *Pc*UndB, for a total of 1.8μs. Combining all these simulations, a total of 1.8 microseconds (μs) of all-atom simulations were performed using a timestep of 1fs, with trajectories recorded every 500 fs for detailed temporal resolution. Long-range electrostatic effects were modelled using the Particle-Mesh Ewald (PME)[7] method. The system temperature and pressure were maintained at 300K and 1bar using the Langevin piston Nóse–Hoover method[8,9], respectively. A 12 Å cutoff was applied to Lennard-Jones and short-range electrostatic interactions. Trajectory analysis was performed using the VMD[4] program. Molecular dynamics (MD) simulations were performed using the GPU-accelerated version of the NAMD[10] package.

### 1.2 QM/MM MD Simulations

We applied weak restraints throughout the production run to maintain the active site in quasi-reactive conformation. Under this condition, no significant conformational changes were observed in the protein structure. Thus, to promote chemically relevant bond reorganization events, we employed a variant of steered molecular dynamics (SMD) simulations called conditional SMD QM/MM MD (c-SMD-QM/MM MD)[11], implemented in the NVT ensemble for 15 ps with a 1 fs integration time step. The simulations used a hybrid QM/MM interface, and atoms within the QM region were biased to induce bond formation or breaking. The c-SMD QM/MM MD steers the simulation within the QM region, while avoiding bringing atoms too close to each other and destabilizing the molecules. A moving harmonic restraint was applied along the reaction coordinate defined as the distance between the activated oxygen and the β-hydrogen of the lauric acid (C12) substrate. The restraint had a force constant of 200.0 $kcal \cdot mol^{-1} \cdot Å^{-2}$ and was advanced at a constant rate of −0.002 $Å.fs^{-1}$. The force constant was chosen to maintain a tight constraint on the system while allowing smooth progression along the reaction coordinate, and the pulling rate was selected to ensure quasi-equilibrium conditions and minimize non-equilibrium artifacts. The force on the atom pairs stops acting once they are closer to the cutoff distance; the simulation continues as unbiased QM/MM MD. This method efficiently explores the reaction pathway. The c-SMD QM/MM MD simulations was followed by 100 ps of unbiased QM/MM MD run. The system was subjected to initial MM minimization using a conjugate gradient algorithm followed by sequential QM/MM minimizations and equilibration. The VMD-QwikMD interface was used for system set up[12]. The QM region consists of the diiron center ($Fe^{2+}$), active site histidines (His89, His93, His128, His131, His132, His261, His297, His300, and His301)[13], the Lauric acid substrate, and water molecules within 2.5Å of the active site. To maintain the charge of the QM region between +1 and -1 for effective semi-empirical calculation[14] some extra residues (Asp90, Glu142, and Glu143) were added to the QM region. The QM region consists of 312 atoms, and the MM layer was composed of the rest of the protein, lipid bilayer, water beyond 2.5 Å, and ions. The PM6-D3H4[15,16] method was employed for the QM region using the NAMD-MOPAC interface [11], while the MM subsystem was treated with CHARMM36 force field[6]. The system was subjected to initial MM minimization using a conjugate gradient algorithm followed by sequential QM/MM minimizations and equilibration through the QwikMD interface[12]. Electrostatic embedding was employed and covalent boundary treatment between QM and MM regions utilized 'link' hydrogen atoms. The total charge and spin multiplicity were set to +1 and 9, 11 respectively. The system was equilibrated under NVT conditions at 300 K for 1ps using a 1 fs integration time step. Temperature control was achieved via a Langevin thermostat, and long-range electrostatics were treated with the particle mesh Ewald (PME) method with a 12 Å cutoff.

### 1.3 QM/MM Free Energy Simulations

The equilibrated intermediate structure from c-SMD QM/MM MD was used as an initial structure for QM/MM well-tempered metadynamics (WT-MetaD)[14,17] simulations to enhance sampling along the collective variable space. QM/MM WT-MetaD simulations were performed at 300 K, 1 bar, 1 fs time step of integration, and periodic boundary conditions for 100 ps using the NAMD-MOPAC [11] interface and the collective variables module. The distance between the β-hydrogen of the substrate and the activated oxygen species, and the distance between the two oxygen atoms were used as collective variables (CVs). Gaussians of height 0.3 kcal/mol were added onto the CV coordinate 10 fs to construct a metadynamics bias potential with a width of 1 Å.QM/MM MD production run using the PM6-D3H4[15] method. The bias potential was tempered using a bias temperature of 1000 K, corresponding to a bias factor of approximately 3.33 at a simulation temperature of 300 K. This scaling gradually reduces the height of successive Gaussians, ensuring smooth exploration of the free energy surface while preventing overfilling. All other simulation parameters were identical to those used in classical QM/MM molecular dynamics.

#### 1.4 QM/MM metadynamics for Fe-Fe distance variation

The initial structure for metadynamics[18,19] simulation was taken from the Alphafold3 UndB after equilibration in ligand-unbound conditions. QM/MM WT-MetaD simulations were performed at 300 K, 1 bar, 1 fs time step of integration, and periodic boundary conditions for 35 ps using the NAMD-MOPAC [11] interface and the collective variables module. The distance between two iron atoms was used as a collective variable (CV). Gaussians of height 0.2 kcal/mol were added onto the CV coordinate 10 fs to construct a metadynamics bias potential with a width of 1 Å. Efforts to obtain an oxo or hydroxo-bridged structure didn't succeed after several attempts.

#### 1.5 Metadynamics for inactive to active conformational transition

The initial structure for metadynamics[18,19] simulation was taken from the lauric acid bound UndB after equilibration. The metadynamics simulation aimed to generate a free energy landscape from the MD-equilibrated structure to the near-attack conformation obtained from c-SMD QM/MM MD. The collective variable (CV) used for the free energy landscape generation was the distance between the oxygen atom ($O_2$) and the Cβ atom of the lauric acid backbone and the distance between Arg124 and Glu143. Two individual metadynamics simulation with the CV distance between the oxygen atom ($O_2$) and the Cβ atom of the lauric acid backbone and the distance between Arg124, and distance between the oxygen atom ($O_2$) and the Cβ atom of the lauric acid backbone and the distance between Glu143 were also ran to understand the individual role of each of these residues. A hill width of 0.01 kcal/mol was used throughout the simulation, which was run for 50 ns each. A total of 150 ns of metadynamics simulation was run.

#### 1.6 Quantum chemical (QM) calculations

We constructed a cluster model of UndB from the appropriate QM/MM MD snapshot. All Quantum chemical calculations were carried out using the ORCA package[20,21]. The electronic structures and vibrational properties of the di-iron active-site model were investigated using a quantum chemical (QM) cluster model approach**.** The cluster model included both Fe (II) centers, their directly coordinating histidine residues necessary to reproduce the local geometry and electronic environment of the enzyme active site. The QM region consists of the diiron center ($Fe^{2+}$), active site histidines (His89, His93, His128, His131, His132, His261, His297, His300, and His301), the lauric acid substrate truncated at the $C_\delta$ position, and a water molecule. Selected peripheral $C_\alpha$ atoms were held fixed during optimization to maintain the constraints imposed by the protein's secondary and tertiary structure, and is selected as a model for full QM calculations. The use of cluster model approach has been pioneered by Siegbahn and coworkers along with others in studying metalloenzymes and it provides accurate results as per other computational methods[22]. The cluster model had 158 atoms in the absence of external proton, finally 159 atoms after adding external proton and electron. The Grimme's dispersion corrected unrestricted the B3LYP[23] hybrid exchange-correlation functional was employed for geometry optimization. Geometry optimization and harmonic frequency calculations were performed for all entities, which include reactants, intermediates, transition states, and products. Additionally, we used the def2-SVP[24] basis set in all the calculations. All optimizations were performed in the presence of a polarizable continuum model (CPCM) (24) to mimic the low-dielectric environment and approximate the electrostatic screening typical of enzyme interiors. The calculations were performed using the unrestricted Kohn-Sham formalism [25] for an overall singlet spin state (S = 0), employing a broken-spin symmetry approach corresponding to the anti-ferromagnetically coupled Fe (II)–Fe (II) configuration of the resting di-iron center. Vibrational frequency analyses were performed in conjunction with geometry optimizations to confirm that each structure corresponds to a true minimum on the potential-energy surface. To locate an appropriate transition-state geometry, a relaxed potential energy surface (PES) scan was performed along the key reaction coordinate. At each point of the scan, the selected coordinate was constrained while all other degrees of freedom were fully optimized at the B3LYP-D3/def2-SVP level of theory. The resulting energy profile revealed a single energy maximum corresponding to the transition state region. Previous studies have confirmed that the transition states (TS) obtained from full optimization and PES scanning only exhibit minor differences in energy and structure[26,27]. We have performed frequency analysis for all the transition states. TS1 exhibits an imaginary frequency of 588.51*i* $cm^{-1}$ corresponding to vibrational mode of $O_A$-$O_B$ cleavage. TS2 exhibits an imaginary frequency of 293.67*i* $cm^{-1}$ involving abstraction of β-H by $O_A$ of Fe-oxo intermediate. TS3 exhibits an imaginary frequency of 209.75*i* $cm^{-1}$ involving cleavage of C1-C2 bond and $CO_2$ release. The results of proper imaginary mode indicates that the structures of transition states are reliable.

**2.0 Structural integrity of the AlphaFold3 structure of UndB**

The 3D structure of UndB was predicted using AlphaFold3. The structure corresponds to UndB from Pseudomonas canadensis [GenBank: UKZ44198.1] The model confidence (pLDDT scores): 90 (very high confidence), core transmembrane helices and structured domains,70–90 (confident), surface helices and amphipathic segments, <70 (low confidence), flexible loops and termini. The Ramachandran analysis shows 95.2% residues in favoured regions,99.2% residues in allowed regions (Fig. S3). Rotamer quality shows 98.99% favored rotamers. Geometric quality shows 0 bad bonds, 0 Cβ deviations, no clashes. Membrane topology prediction shows transmembrane helices (TM1–TM4), Cytosolic N-terminus, extracellular C-terminus, Amphipathic and soluble helices flank the membrane core. Matches well with structural prediction from AlphaFold. Based on the averaged predictions from six topology tools, the protein is most reliably predicted to contain four transmembrane helices: TM1 spanning residues 42 to 60, TM2 from 63 to 80, TM3 from 187 to 207, and TM4 from 230 to 250.

(a)

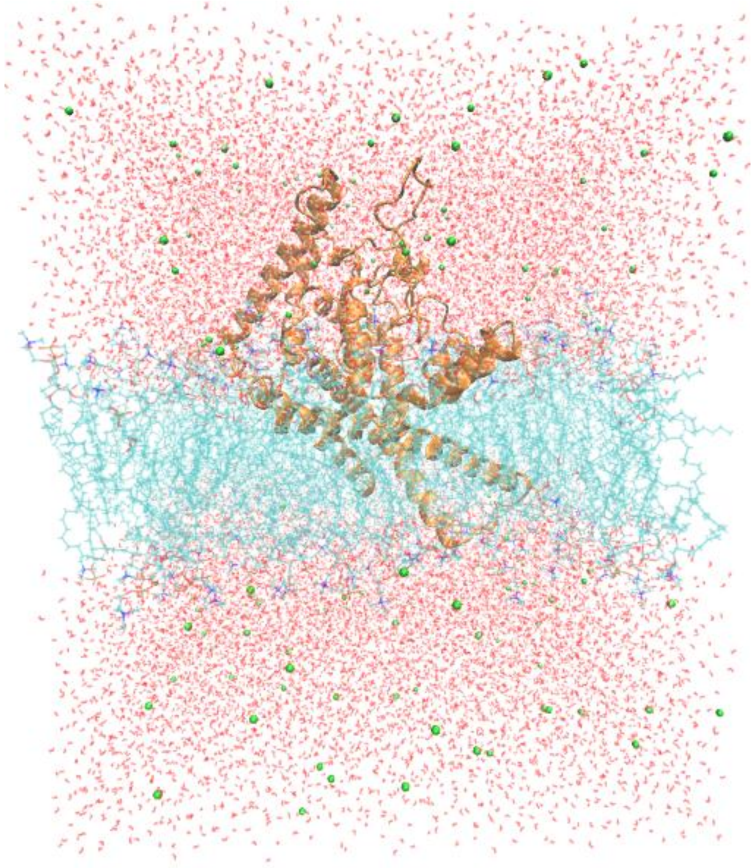

(b)

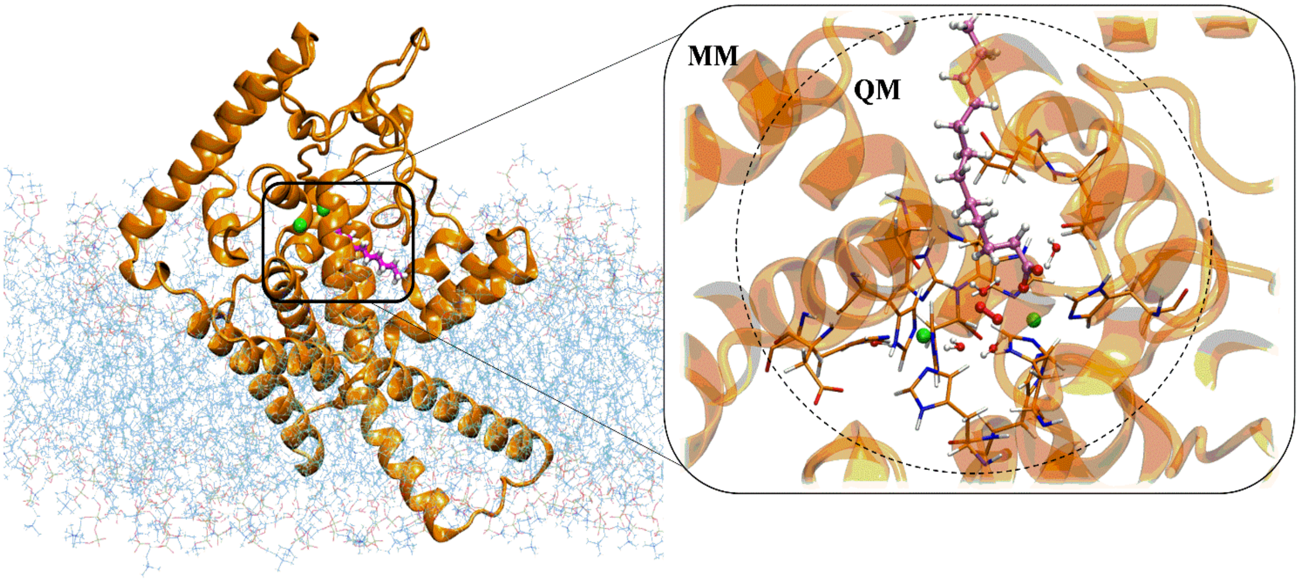

**Fig. S1.** Setup of the molecular simulations of UndB. (a) Setup of the classical MD simulations. (b) Setup of the QM/MM calculations (left), and a closeup of the 312 atoms included in the QM region of QM/MM MD simulations (inset, right).

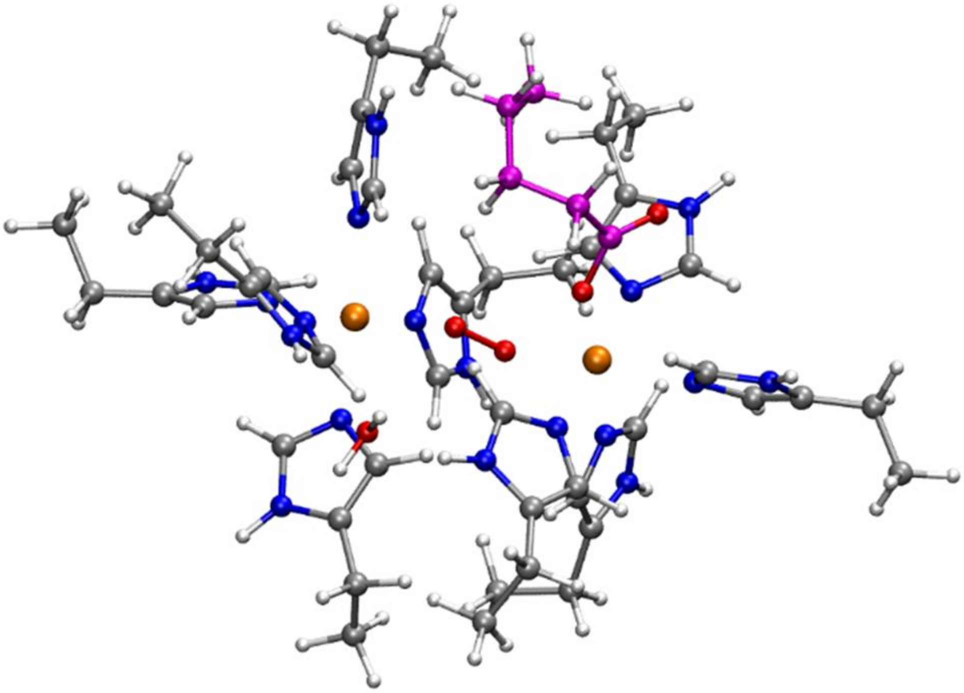

**Fig. S2.** The QM region selected for high-level cluster model quantum chemical calculations at the B3LYP-D3/def2-SVP level of theory.

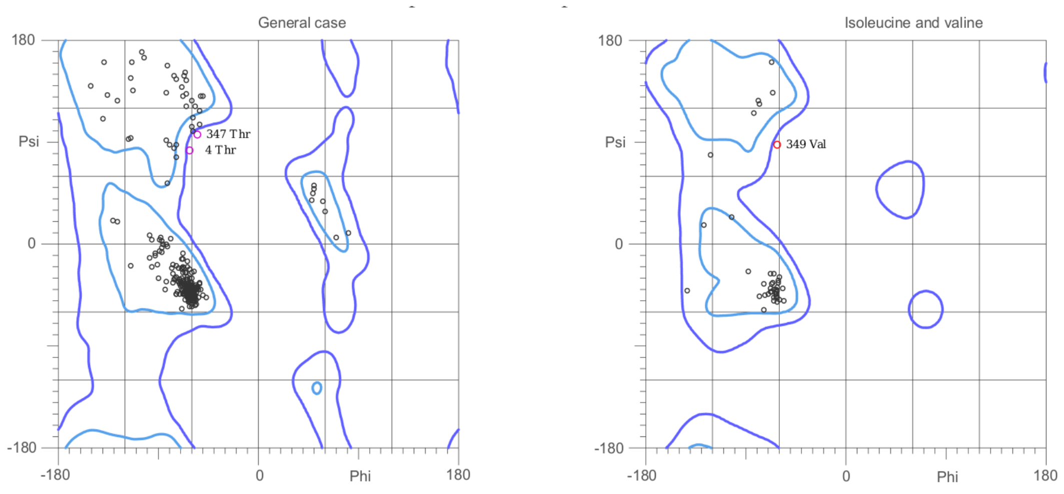

**Fig S3.** Ramachandran plot of UndB.

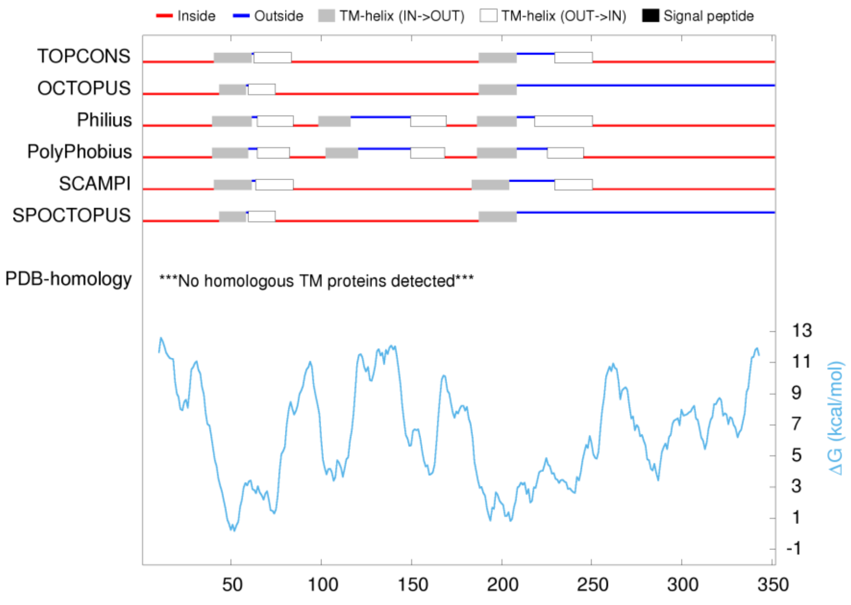

**Fig S4.** Transmembrane topology of UndB.

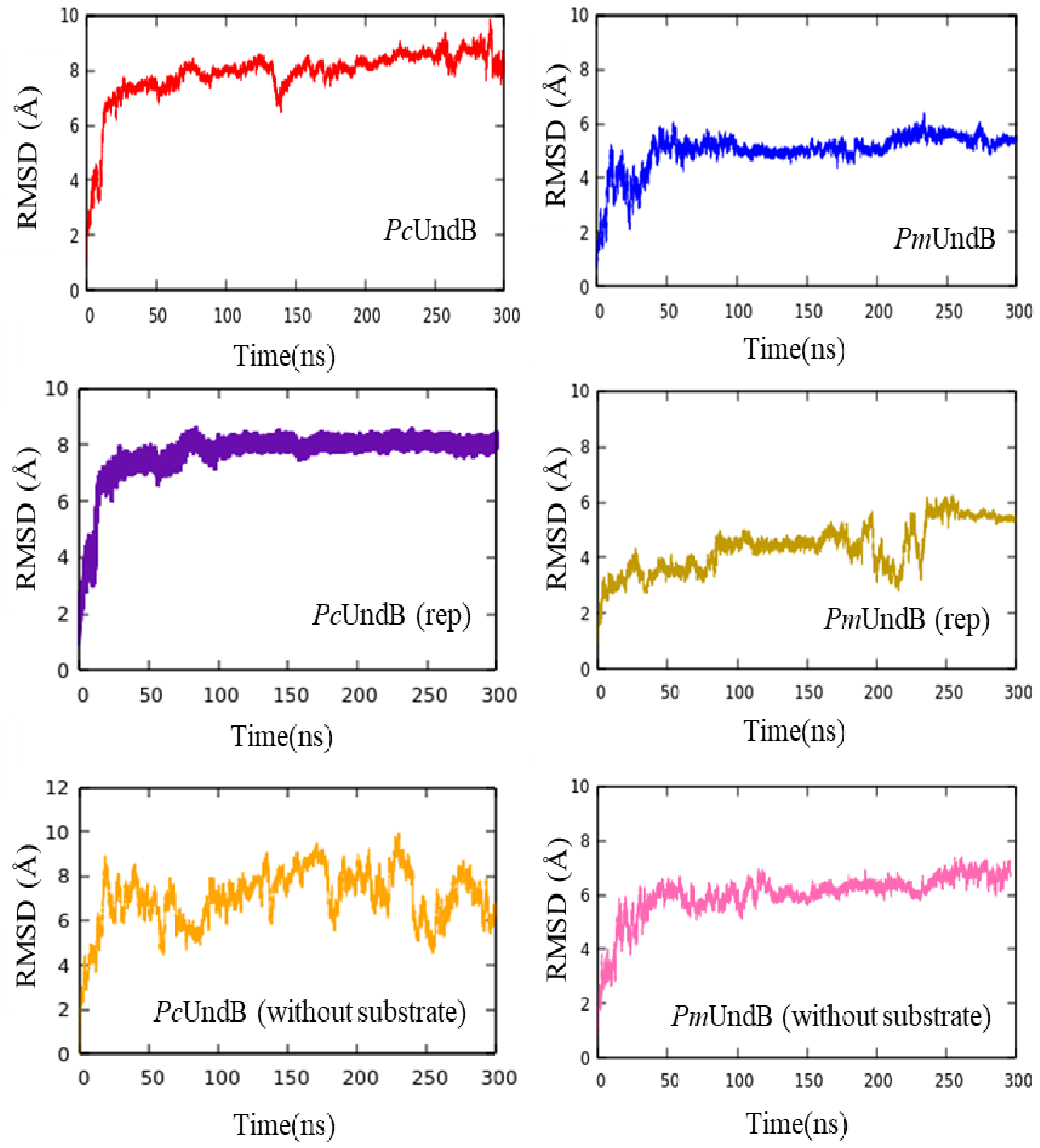

**Fig. S5.** Protein backbone ($C_\alpha$) Root Mean Square Deviation (RMSD) plots for two UndB structures and its replicas from a total 1.8 μs all-atom MD simulations trajectory. (rep) refers to replica of the corresponding UndB structure.

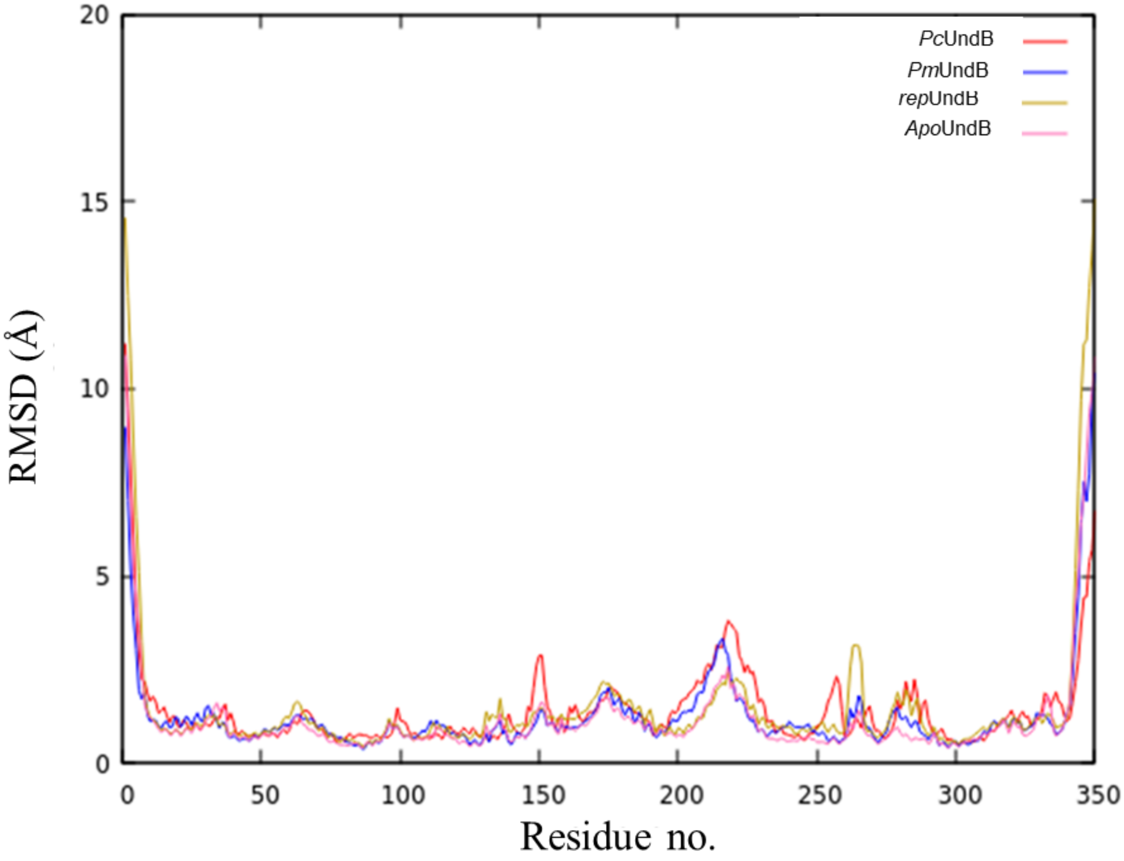

**Fig. S6.** RMSF plots for the protein backbone ($C_\alpha$) atoms for 300 ns of the simulations, each shown for three substrate-bound UndB and apo-UndB structures.

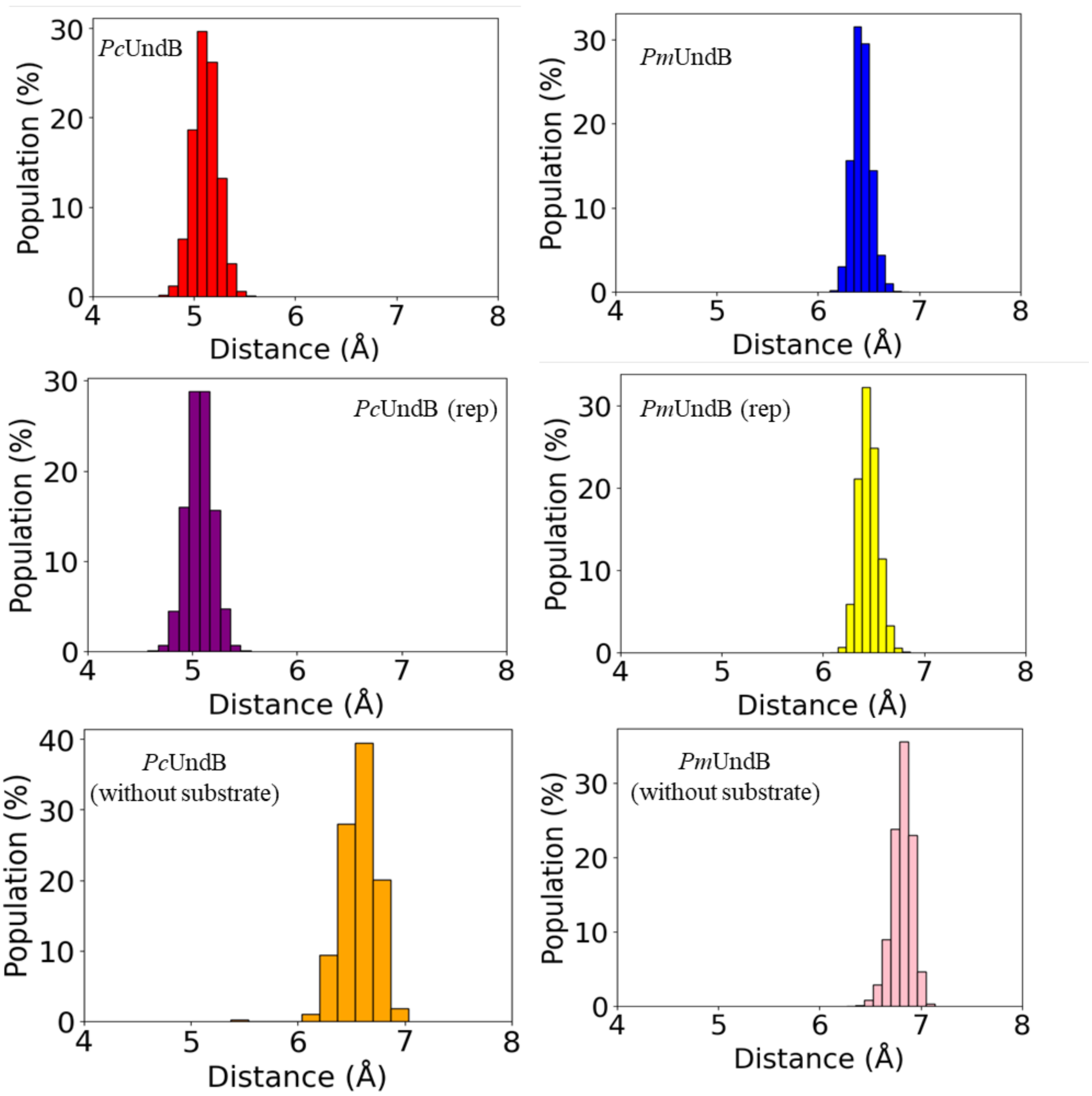

**Fig. S7.** Distribution of Fe…Fe distances from classical molecular dynamics simulations of replicas of substrate-bound UndB and substrate-unbound UndB. (rep) refers to replica of the corresponding UndB structure.

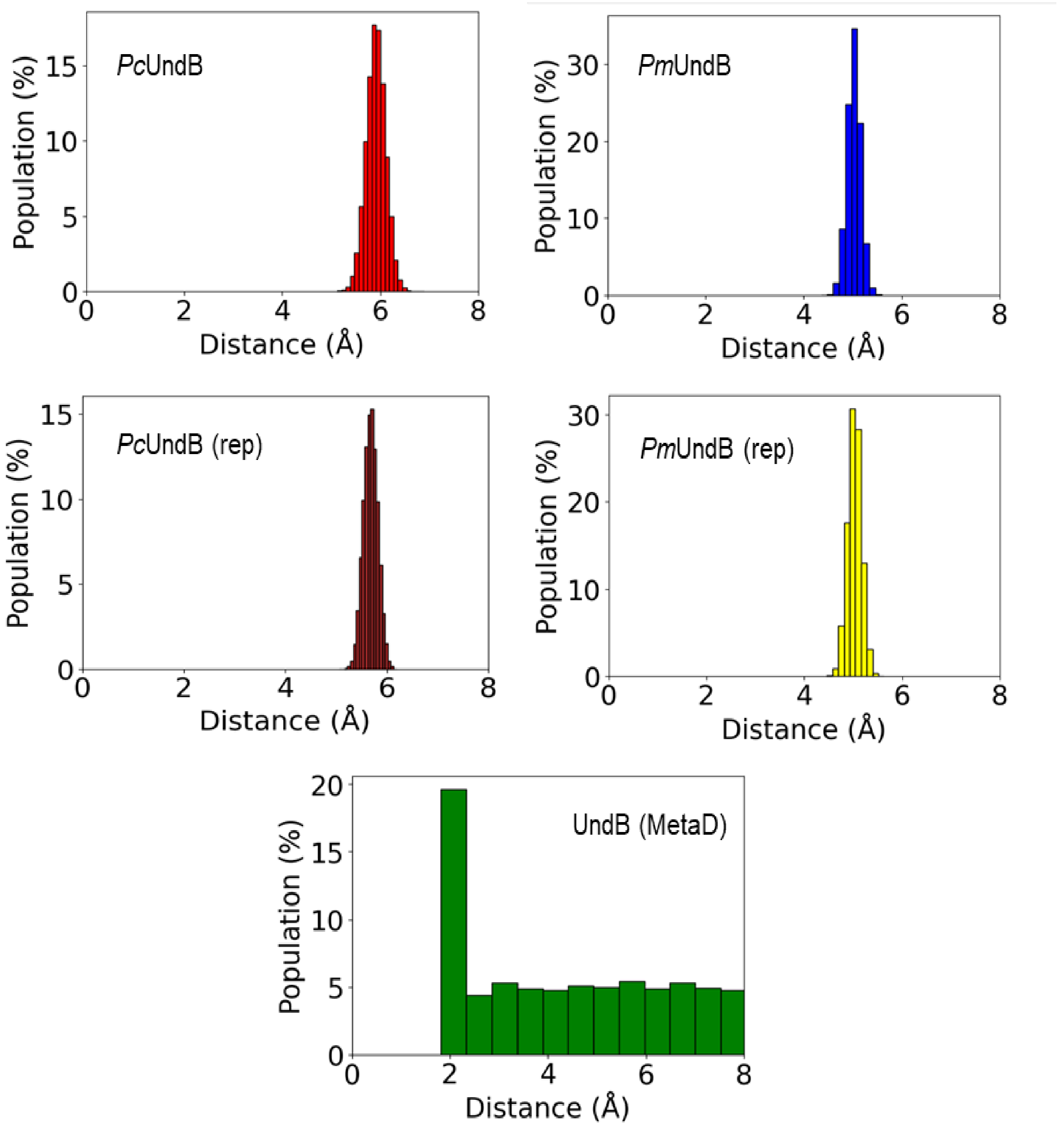

**Fig. S8.** Distribution of $O_B$ ($O_2$)…β-H distances for different replicas of substrate-bound UndB. MetaD refers to classical metadynamics simulations. (rep) refers to replica of the corresponding UndB structure.

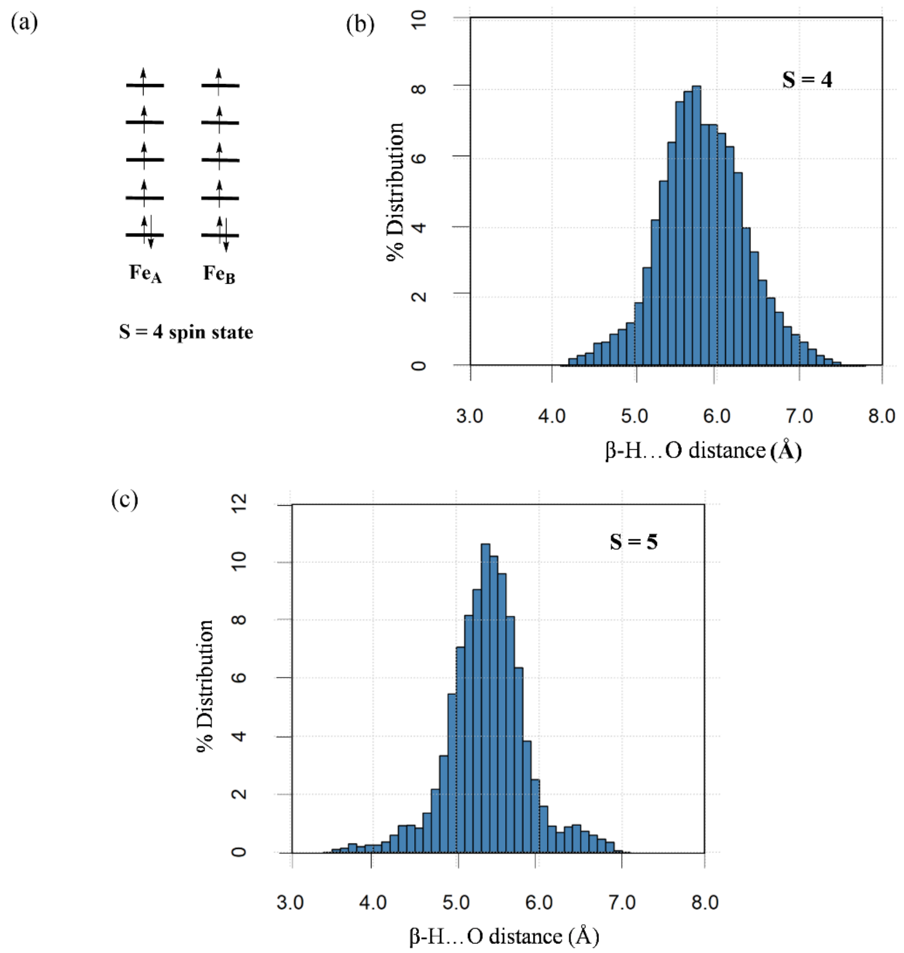

**Fig. S9.** (a) Schematic representation of the high spin states (S = 4 and 5) of the $ESO_2$ complex of UndB considered in the QM/MM MD simulations. Distribution of the $O_A(O_2)$…β-H distances for (b) high spin S = 4 state and (c) high spin S = 5 state from QM/MM MD simulations of the substrate-bound UndB.

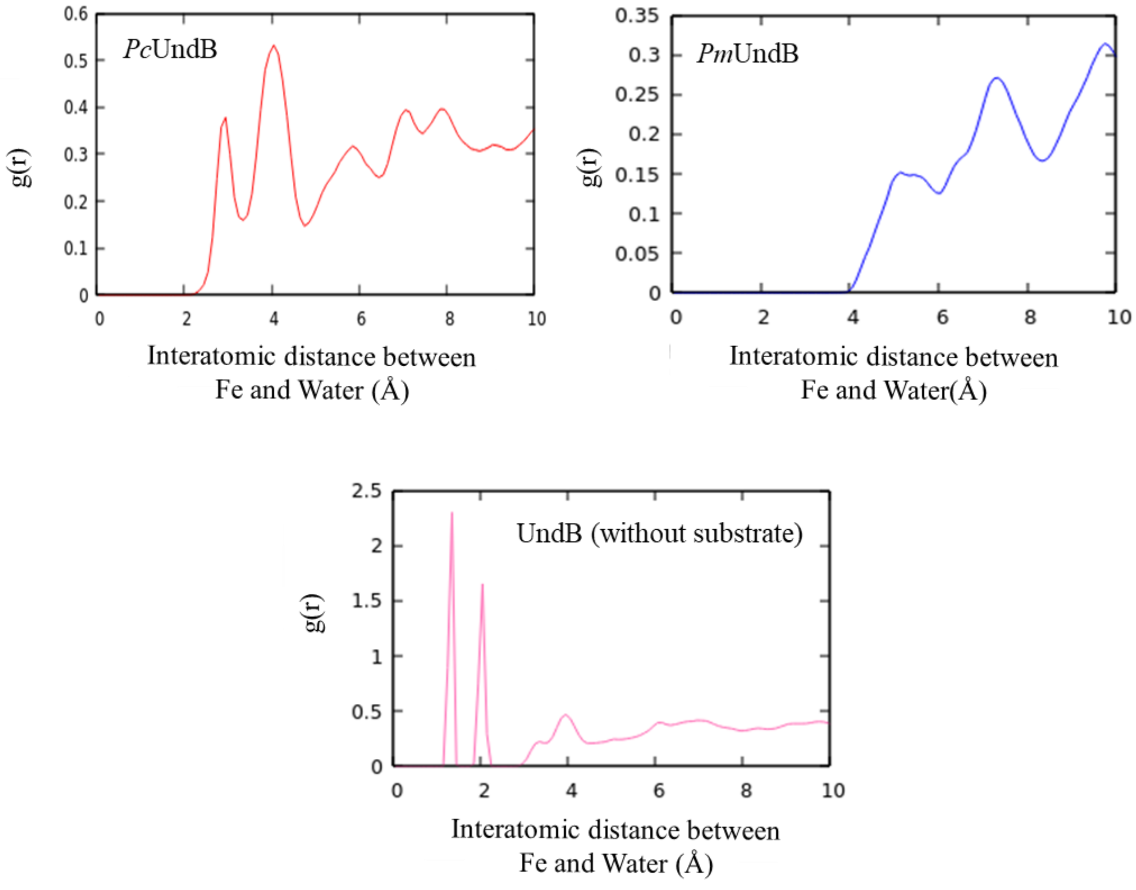

**Fig. S10.** Radial Distribution Function (RDF) plots measured from different UndB simulation trajectories.

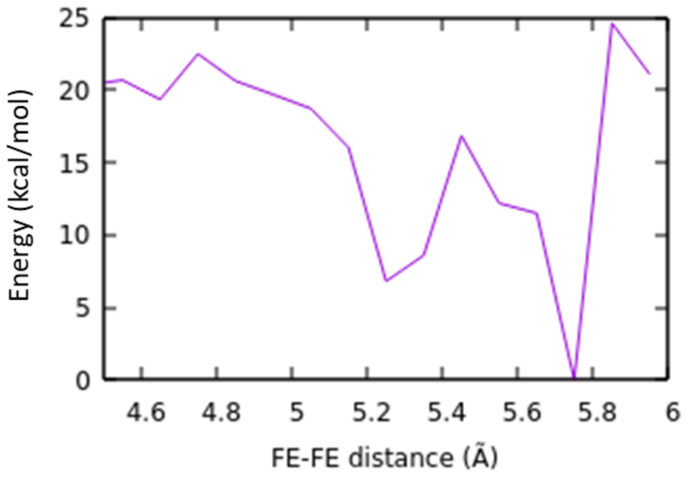

**Fig. S11.** QM/MM metadynamics simulations along the Fe-Fe distance.

#### 3.0 Nature of the reactive oxygen species that performs β-H abstraction in UndB

We optimized the peroxo-like intermediate **(P)** obtained from QMMM MD simulations using the cluster model (**P1**) at the open-shell singlet state (S = 0) and the nonet spin state (S = 4) along with other relevant spin state (S = 5) at the B3LYP-D3/def2-SVP level of theory. The open-shell singlet is found to be 4.1 kcal/mol lower in energy than the nonet spin state and lies 1.0 kcal/mol lower than the highest spin state and is the ground state. The highest spin state (S = 5) doesn't show any β-H abstraction from the substrate in QM/MM MD simulations. This may be attributed to the $O_2$ being not activated in the highest spin state as clear from the calculated QM spin densities on the $O_A$ (0.50) and $O_B$ (0.46) atoms[28,29]Notably, $O_2$ activation occurs only in the presence of substrate (lauric acid) coordinating to the $Fe_B$ center (Tables S1 and S2). $O_2$ binding to the LA-coordinated $Fe_B$ site results in the formation of a μ-1,2-peroxodiferric complex, which is accompanied by contraction of the Fe–Fe distance from 5.8 Å to 4.9 Å (Fig S12a). In this context, it is worth recalling that the first crystallographic structure of UndA in complex with a substrate analogue reported a trapped superoxo species[30]; however, subsequent characterization unambiguously reassigned this species a peroxodirron(III/III) species using Mossbäuer spectroscopy[30]. However, from QM calculations we found that the formed μ-1,2-peroxodiiron(III/III) intermediate **(P1)** is kinetically incapable of promoting substrate activation. The calculated barriers for β-H abstraction from lauric acid (LA) are prohibitively high, amounting to 62.8 kcal/mol and 46.5 kcal/mol for hydrogen abstraction by $O_A$ and $O_B$, respectively. In addition, cleavage of the $O_A$–$O_B$ bond requires 25.6 kcal/mol, leading to formation of the corresponding bis-Fe=O species **(FO2)** from the peroxodiiron(III/III) complex that also does not resolve this limitation. The resulting diiron-oxo intermediate **(FO2)** exhibits β-H abstraction barriers of 32.0 kcal/mol (by $O_A$) and 53.0 kcal/mol (by $O_B$), values that remain incompatible with a catalytically viable pathway.

Hence, we optimize the hydroperoxo intermediate **(IM1)** by adding an external proton and an electron to the peroxodirron(III/III) species (Fig. S12 b,c). Upon protonation of the $O_B$ of the peroxo-bridge, the reaction proceeds through **TS1** with a barrier of 7.5 kcal/mol to the iron-oxo intermediate **(IM2)**. The $Fe_A$=$O_A$ intermediate abstracts the β-H with a barrier of 15.0 kcal/mol **(TS2)** to form a substrate radical intermediate **(IM3)**, in match with the experimentally observed barrier of 17.9 kcal/mol. Proton transfer from $O_B$ to $O_A$ led to the formation of intermediate **(IM4)**. The final step was carbon dioxide release, which proceeds through a barrier height of 4.3 kcal/mol **(TS3)** to form the product, the 1-alkene. We also tried by adding only a proton; in the absence of an external electron, the intermediate gave a barrier of 28.0 kcal/mol for β-H abstraction if $O_A$ abstracts and 40.0 kcal/mol if $O_B$ abstracts. Thus, it is evident that supply of external and electrons and protons is essential for UndB catalysis.

#### 3.1 Alternative $O_2$ activation pathways

We further examined whether alternative $O_2$ activation pathways could be operational in UndB. To this end, we evaluated superoxo-like intermediate formation by coordinating $O_2$ to $Fe_A$ and $Fe_B$ separately in both the open-shell singlet and nonet spin states (Fig. S13). In both cases, these Fe-bound superoxo-like species were energetically disfavoured relative to the ground-state peroxodiiron(III/III) intermediate **(P1)**. Specifically, the $Fe_A$-bound superoxo **(S1)** lies 11.7 kcal/mol (high-spin) and 12.6 kcal/mol (open-shell singlet) above the peroxo intermediate, whereas the corresponding $Fe_B$-bound superoxo **(S2)** is higher in energy by 6.4 kcal/mol in both

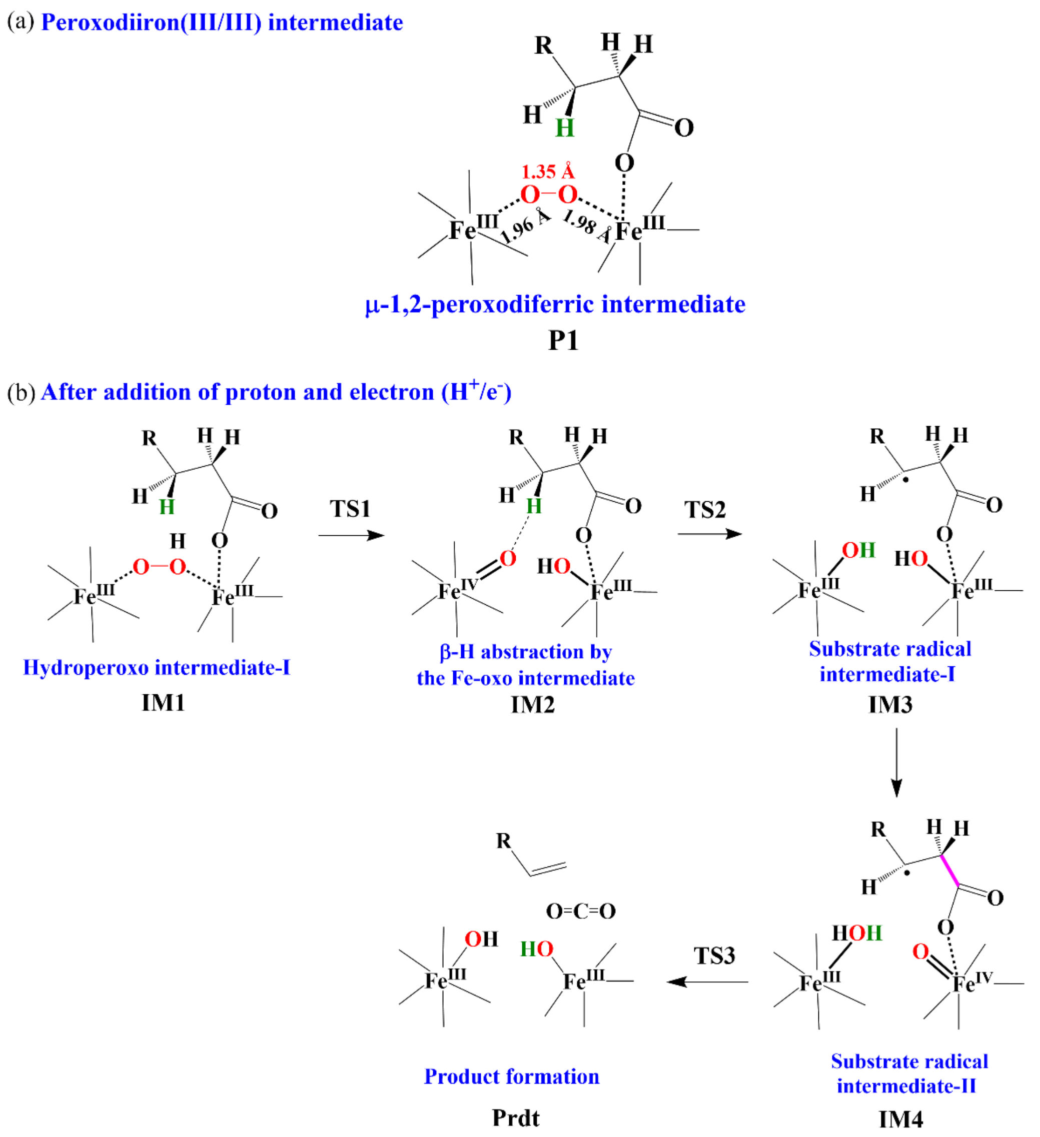

**Fig S12.** (a) Schematic representation of the optimized peroxodiiron(III/III) intermediate from cluster model quantum chemical calculations at the B3LYP-D3/def2-SVP level of theory. (b) Proposed catalytic pathway of UndB through the formation of a (hydro)peroxo intermediate at the B3LYP-D3/def2-SVP level of theory.

spin states, also $O_A$ which is supposed to abstract β-H is not activated as well as oriented distantly in **S2**. We next explore the protonation pathways by considering the active-site water molecule as a potential proton donor. Geometry optimization of the resulting hydroperoxo species at the nonet spin state yielded intermediates that remain higher in energy, lying 15.9 kcal/mol **(HP1)** and 18.6 kcal/mol **(HP2)** (Fig. S13) above the peroxo reference state. Finally, to emulate formation of a hydroperoxo intermediate under reductive conditions, we introduced an external proton and electron, generating the corresponding S = 4.5 state for further analysis. This hydroperoxo intermediate lies 0.8 kcal/mol lower in energy compared to the hydroperoxo reference state (**IM1**), when –OOH is coordinated to $Fe_A$ **(HP3)** and 1.9 kcal/mol higher when bound to $Fe_B$ **(HP4)**. Despite this apparent thermodynamic viability, cleavage of the $O_A$–$O_B$ bond in the hydroperoxo model requires a substantial barrier of 17.4 kcal/mol. The magnitude of this barrier renders the pathway kinetically higher, leading us to exclude this mechanistic scenario.

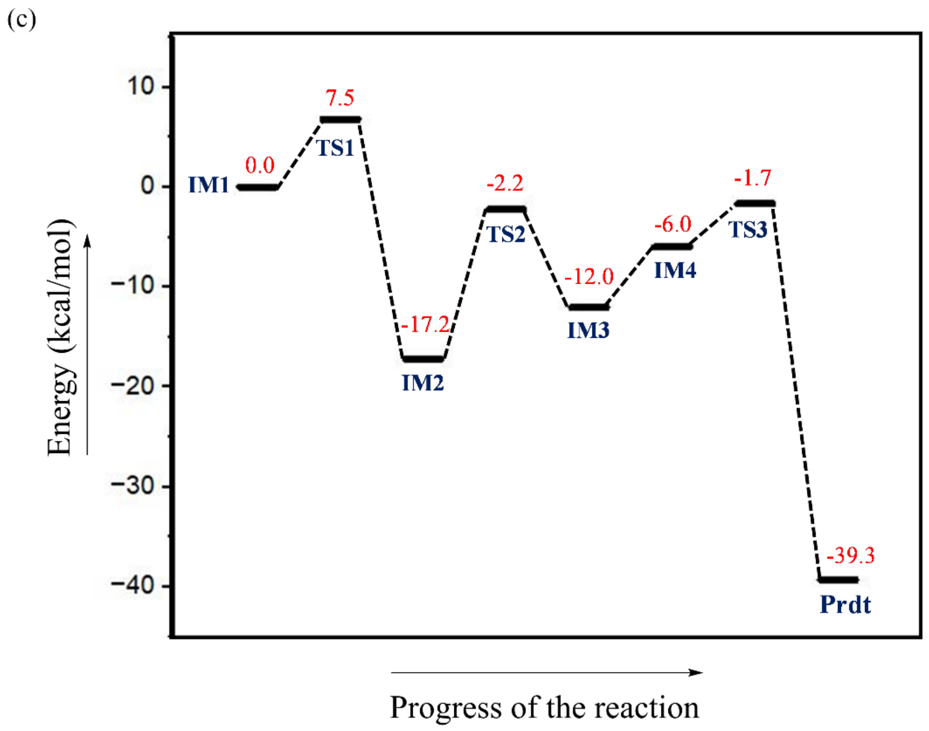

**Fig S12.** (c) Reaction energy profile of the catalytic pathway of UndB obtained using quantum chemical cluster model calculations at the B3LYP-D3/def2-SVP level of theory.

To further substantiate the proposed mechanism, we systematically examined a range of alternative oxygen-derived intermediates. Attempts to optimize **S1** and **S2** (Fig. S14) as open-shell singlets were unsuccessful; in both cases, geometry optimization spontaneously converged to the peroxo-bridged species **(P1)**. **P2** similarly relaxed to a structure closely resembling **P1**, underscoring the intrinsic stability of the peroxo-bridged motif within the UndB active site. **HP5** could be optimized at the S = 4.5 spin state, where the high-spin configuration was marginally favoured over the broken-symmetry solution by 0.3 kcal/mol; however it was found have a barrier of 28.4 kcal/mol for subsequent O-O bond breaking step. **FO1** was likewise optimized at S = 4.5, with the high-spin state stabilized by 5.0 kcal/mol relative to the broken-symmetry state, and showed higher barrier of 32.5 kcal/mol for β-H abstraction. In contrast, **HP6** could not be retained during optimization; instead, it evolved into an unbridged structure featuring –OOH bound to $Fe_B$ and a hydroxide ligand coordinated to $Fe_A$. Formation of a high-valent Q-like species **(Q)** resulted in an unbridged bis-Fe(IV)=O **(FO2)** motif upon optimization. Although **FO2** could be located as a stationary point, it lies substantially above the ground state (15.2 kcal/mol in the high-spin state and 15.4 kcal/mol in the broken-symmetry state). Attempts to generate **FO3** instead led to relaxation into the same unbridged **FO2** configuration. Moreover, even though **FO2** could be stabilized, the computed β-H abstraction barriers were prohibitively high (32.0 kcal/mol for $O_A$ and 42.0 kcal/mol for $O_B$ being the abstracting species), rendering this pathway kinetically implausible. Collectively, these results reinforce the mechanistic preference for the peroxodiiron(III/III) intermediate and disfavour alternative superoxo, hydroperoxo, and high-valent bis-oxo intermediates.

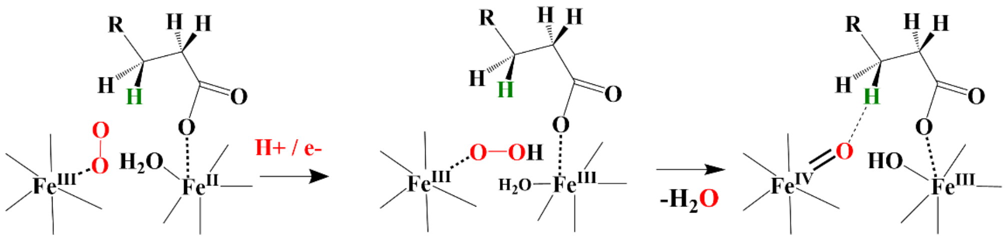

**Fig S13.** Possible alternative catalytic pathway involving different roles of the two iron centers in UndB at the B3LYP-D3/def2-SVP level of theory.

**Table S1.** Important structural parameters of the optimized structures of the intermediates involved in the catalytic reaction pathway of UndB at the B3LYP-D3/def2-SVP level of theory.

| **Structural Parameters** | **Peroxo (P1)** | **Hydroperoxo (IM1)** | **Fe-oxo (IM2)** | **Substrate Radical I (IM3)** | **Substrate Radical II (IM4)** | **Product (Prdt)** |
|---|---|---|---|---|---|---|
| d($Fe_A$...$Fe_B$) | 4.86 | 5.40 | 5.64 | 5.65 | 5.69 | 5.60 |
| d($Fe_A$-$O_A$) | 1.96 | 1.93 | 1.86 | 1.84 | 2.22 | 2.07 |
| d($Fe_B$-$O_B$) | 1.98 | 2.48 | 1.63 | 1.86 | 1.63 | 1.76 |
| d($Fe_B$-O(LA)) | 1.95 | 2.05 | 2.03 | 2.02 | 2.03 | 2.85 |
| d($O_A$-$O_B$) | 1.35 | 1.40 | 2.73 | 2.76 | 2.66 | 2.42 |
| d($O_A$...$H_\beta$) | 3.40 | 3.67 | 3.56 | 0.97 | 0.98 | 0.97 |
| d($O_B$...$H_\beta$) | 4.10 | 4.11 | 4.14 | 3.02 | 3.08 | 2.81 |
| d(C1-C2) | 1.52 | 1.53 | 1.52 | 1.53 | 1.54 | 1.34 |

**Table S2.** Spin densities of the optimized structures of all the intermediates involved in the catalytic reaction pathway of UndB at the B3LYP-D3/def2-SVP level of theory.

| **Intermediates** | **Spin Multiplicity** | **Spin density** | | | | | |
|---|---|---|---|---|---|---|---|
| | | $Fe_A$ | $Fe_B$ | $O_A$ | $O_B$ | β-C | O (LA) |
| Peroxo **(P1)** | 1 | -4.08 | 4.21 | -0.37 | 0.07 | 0.00 | 0.15 |
| Peroxo **(P1)** | 9 | 3.82 | 4.26 | -0.50 | -0.27 | 0.00 | 0.17 |
| Hydroperoxo I **(IM1)** | 10 | 4.18 | 3.83 | 0.35 | 0.08 | 0.00 | 0.04 |
| Fe-oxo **(IM2)** | 10 | 3.15 | 4.26 | 0.62 | 0.32 | 0.00 | 0.11 |
| Substrate radical I **(IM3)** | 10 | 2.90 | 4.25 | 0.01 | 0.34 | 1.01 | 0.11 |
| Substrate radical II **(IM4)** | 10 | 3.82 | 3.23 | 0.04 | 0.54 | 0.99 | 0.07 |
| Product **(Prdt)** | 10 | 3.81 | 4.15 | 0.06 | 0.55 | 0.00 | 0.00 |

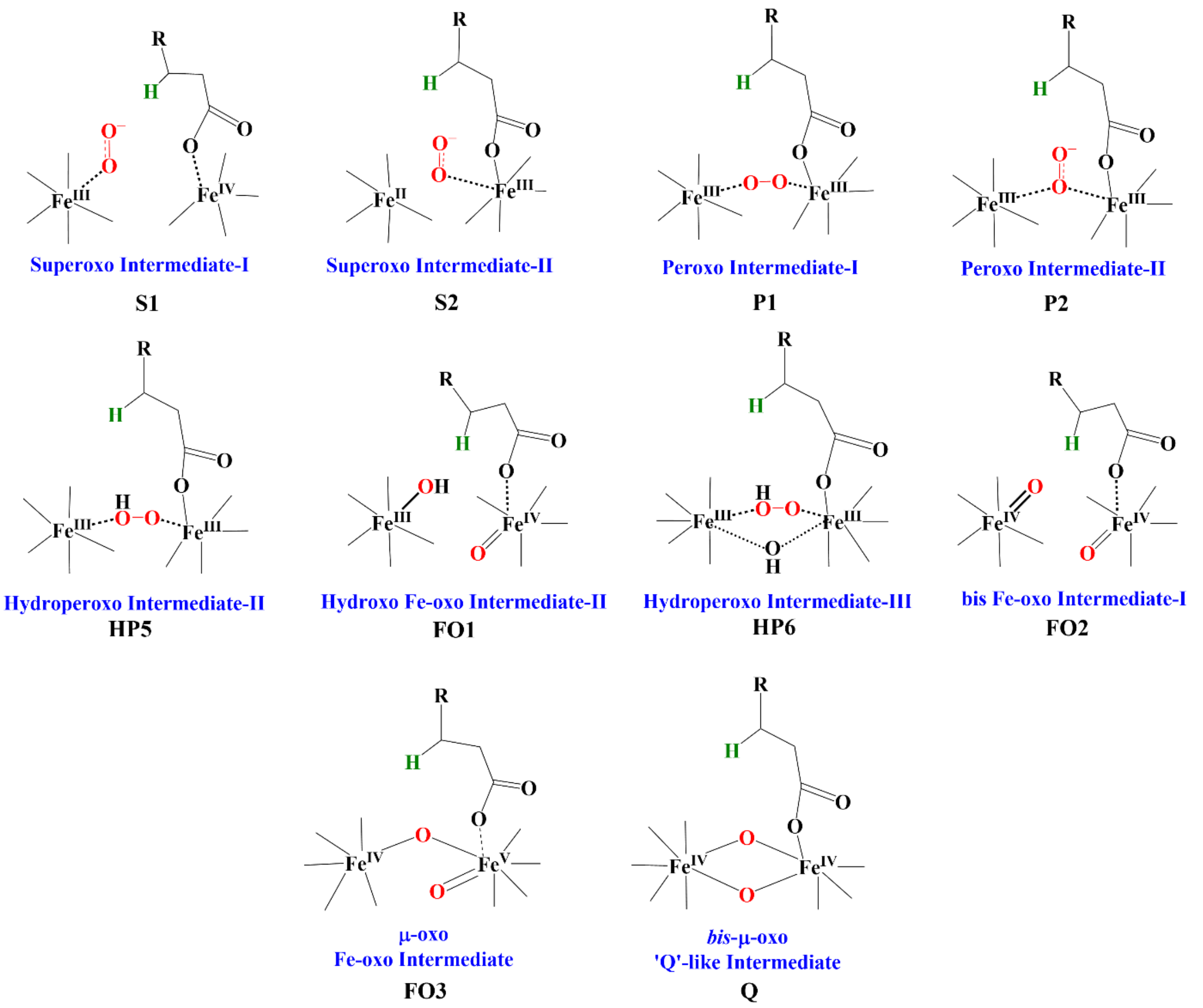

**Fig. S14.** Possible alternative intermediates investigated in this study for likely involvement in the UndB catalytic pathway at the B3LYP-D3/def2-SVP level of theory.

**Table S3.** Important structural parameters of the optimized structures of other alternative intermediates that may be involved in the catalytic reaction pathway of UndB at the B3LYP-D3/def2-SVP level of theory.

| **Structural Parameters** | **Superoxo ($Fe_A$) (S1)** | **Superoxo ($Fe_B$) (S2)** | **Hydroperoxo ($Fe_A$) (HP1)** | **Hydroperoxo ($Fe_B$) (HP2)** | **Hydroperoxo ($Fe_A$) with $H^+/e^-$ (HP3)** | **Hydroperoxo ($Fe_B$) with $H^+/e^-$ (HP4)** |
|---|---|---|---|---|---|---|
| d($Fe_A$...$Fe_B$) | | | | | | |
| S=4 | 5.80 | 5.85 | 5.95 | 5.85 | | |
| S=0 | 5.85 | 5.85 | 5.90 | | | |
| S=4.5 | | | | | 5.75 | 5.75 |
| d($Fe_A$-$O_A$) | | | | | | |
| S=4 | 2.16 | 2.23 | 1.82 | 1.93 | | |
| S=0 | 2.15 | 2.23 | 1.93 | | | |
| S=4.5 | | | | | 1.93 | 2.19 |
| d($Fe_B$-$O_B$) | | | | | | |
| S=4 | 2.07 | 2.05 | 1.87 | 1.81 | | |
| S=0 | 2.06 | 2.05 | 1.87 | | | |
| S=4.5 | | | | | 2.26 | 1.98 |
| d($O_A$-$O_B$) | | | | | | |
| S=4 | 1.32 | 1.29 | 1.43 | 1.43 | | |
| S=0 | 1.32 | 1.29 | 1.41 | | | |
| S=4.5 | | | | | 1.42 | 1.42 |

**Table S4.** Spin densities of the optimized structures of other alternative intermediates that may be involved in the catalytic reaction pathway of UndB at the B3LYP-D3/def2-SVP level of theory.

| **Intermediates** | **Spin Multiplicity** | **Spin Density** | | | |
|---|---|---|---|---|---|
| | | **$Fe_A$** | **$Fe_B$** | **$O_A$** | **$O_B$** |
| Superoxo ($Fe_A$) **(S1)** | 9 | 3.77 | 4.29 | -0.31 | -0.62 |
| | 1 | -3.87 | 4.29 | -0.37 | -0.61 |
| Superoxo ($Fe_B$) **(S2)** | 9 | 3.82 | 4.25 | -0.19 | -0.60 |
| | 1 | -3.82 | 4.25 | -0.19 | -0.60 |
| Hydroperoxo ($Fe_A$) **(HP1)** | 9 | 2.86 | 4.25 | 0.03 | 0.00 |
| | 1 | -4.20 | 4.26 | -0.32 | -0.06 |
| Hydroperoxo ($Fe_B$) **(HP2)** | 9 | 2.87 | 4.22 | 0.28 | 0.04 |
| Hydroperoxo ($Fe_A$) with $H^+/e^-$ **(HP3)** | 10 | 4.21 | 3.83 | 0.33 | 0.06 |
| Hydroperoxo ($Fe_B$) with $H^+/e^-$ **(HP4)** | 10 | 3.82 | 4.26 | 0.20 | 0.03 |

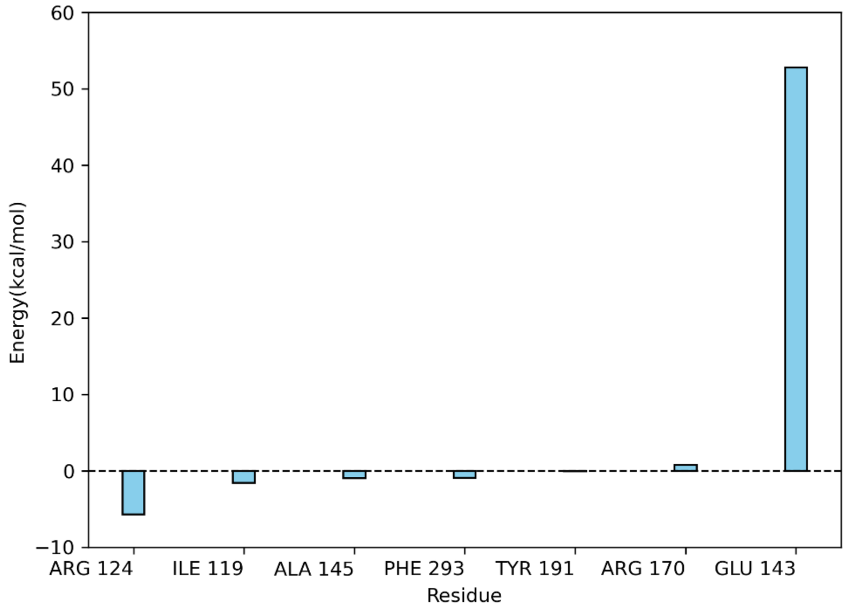

**Fig. S15.** Total non-bonded interactions in active conformation of UndB from classical metadynamics simulations.

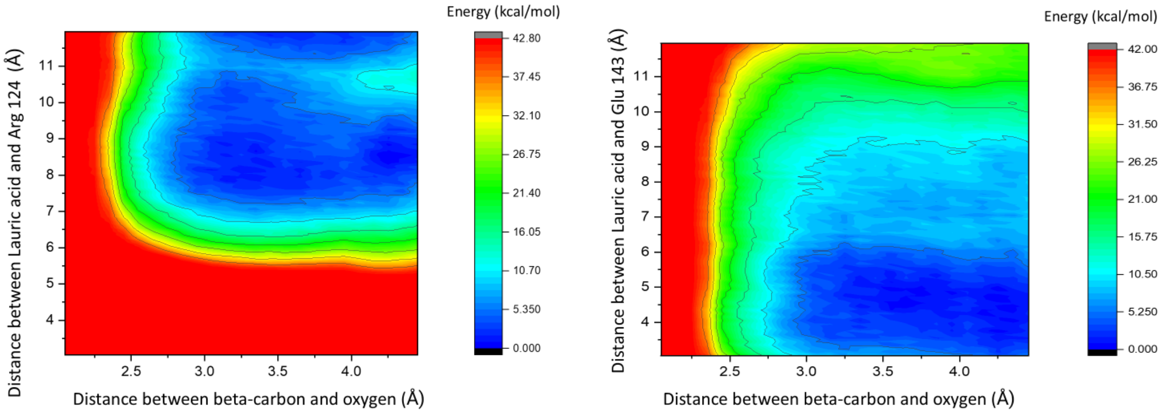

**Fig. S16.** QM/MM metadynamics simulations along the Fe-Fe distance.
